# Both cell autonomous and non-autonomous processes modulate the association between replication timing and mutation rate

**DOI:** 10.1101/2022.10.03.510640

**Authors:** Oriya Vardi-Yaacov, Adar Yaacov, Shai Rosenberg, Itamar Simon

## Abstract

Cancer somatic mutations are the product of multiple mutational and repair processes, both of which are tightly associated with DNA replication. Mutation rates (MR) are known to be higher in late replication timing (RT) regions, but different processes can affect this association. Systematic analysis of the mutational landscape of 2,787 tumors from 32 tumor types revealed that the tumors can be divided into two groups with approximately one third of the tumor samples show weak association between replication timing and mutation rate. Analyses of the two groups revealed that both mutational signatures and mutation in genes involved in DNA replication, DNA repair and chromatin structure impact the association between RT and MR. Surprisingly, analysis of differentially expressed genes between the two groups revealed involvement of cell-cell communication and of the interaction with immune cells in modulating the effect of RT on MR. This provides evidence of the recently described association between mutagenic processes and the immune system in patients’ tumor samples.

## Introduction

The process of DNA replication plays an important role in mutagenesis [1], with failures leading to the introduction of mismatches and/or to the conversion of DNA damages into mutations. It is therefore not surprising that replication timing (RT), defined as the time in S phase each region is replicated, is strongly associated with mutation rate (MR) in both germline and somatic cells. In general, there are many more mutations in regions that replicate in late S phase than in those which replicate in early S phase (reviewed in [2]), suggesting that either mutagenesis, repair or both occur in different intensities in early and late replicating regions.

Mutational signatures are unique combinations of mutations characteristic of various mutagenesis processes [3]. We have recently found that the association between replication timing and mutational signatures differs for different cancer mutational processes [4].

A possible approach for finding the mechanisms that distinguish between mutagenesis in early and late replicating regions (ERR and LRR) is to explore mutation rates in cells harboring a mutation in key genes that are components of the DNA replication or repair mechanisms. Indeed, such an approach was successfully carried out by two groups who found that tumors with defects in either mismatch repair (MMR) or global genome nucleotide excision repair (GG-NER) mechanisms [5, 6] do not show higher mutation rates in LRR, suggesting that differences in the efficiency of DNA repair mechanisms are the basis for the difference in mutation rates between ERR and LRR.

Replication stress is a hallmark of cancer cells that is associated with increased genomic instability. Replication stress is a double-edged sword for cancer cells. While it promotes tumorigenesis by increasing genomic instability, it also hinders their potential to proliferate by destabilizing replication forks [7], sensitizes them to chemotherapy and generates neo-antigens that expose them to immunotherapy [8–10]. This vulnerability is classically exploited in cancer treatment to increase replication stress to unsustainable levels [8, 11], but new strategies are emerging, exploiting recently identified specificities of the replication stress response. Combination approaches integrating replication stress–inducing agents, such as carboplatin or gemcitabine, with immunotherapies like the immune checkpoint inhibitor nivolumab have advanced to clinical trials (NCT02944396, NCT03662074, NCT03061188, NCT02734004, NCT02849496, NCT02657889 and NCT02571725).

In addition to cell autonomous effects of DNA damage, it was recently shown that DNA damage and especially replication stress can recruit the immune system to the damaged cell[12]. It has been proposed that DNA damage and replication stress elicit the activation of inflammatory responses that contribute to tumorigenesis in some contexts and to senescence/aging in others [13, 14]. Recent studies have found that the two systems can have mutual effects on each other. On the one hand, defects in processing DNA replication stalled forks lead to accumulation of cytosolic DNA and to activation of the cGAS–STING pathway, resulting in the activation of the type I IFN pathway with consequent expression of ISG15 (interferon-stimulated gene 1515) [15]. On the other hand, inflammation itself can cause replication stress. A recent study found that high levels of ISG15, intrinsic or induced by interferon-β, accelerates DNA replication fork progression, resulting in extensive DNA damage and chromosomal aberrations [16]. Despite the growing evidence of association between replication stress and the immune system, a direct link between mutagenesis and the immune system in tumor samples has not yet been shown.

Here we readdressed the question of the contribution of RT to mutational distribution by identifying tumors in which this association is weaker. A systematic analysis of the association between RT and mutation rates using 2787 whole-genome sequenced (WGS) tumors, which are available from the Pan-Cancer Analysis of Whole Genomes (PCAWG; [17]), reveals that the association of approximately a third of the samples is much weaker than that of the majority of samples. We hypothesized that analyses of the two groups of tumors, which differ in association between RT and MR, will help us to understand the molecular basis for this association. We show that mutational signatures of DNA repair, DNA replication and chromatin organization pathways all contribute to the identified differences. Interestingly, analysis of genes differentially expressed between these two groups of tumors revealed involvement of cell-cell communication and of the interaction with immune cells in modulating the effect of RT on mutation rates. These findings go along with recent observations about a link between replication stress and immune response and, as far as we know, this is the first such relationship reported in vivo. Taken together, our comprehensive approach reveals the involvement of both known and novel processes in controlling the genomic distribution of cancer somatic mutations.

## Results

### Using the degree of association between mutation rates and replication timing to characterize tumor samples

Mutation rates (MR) are associated with replication timing (RT). As a rule, MR are higher in LRR (late replication regions) and lower in ERR (early replication regions) [2, 4]. However, analysis of individual tumors revealed that this association varies considerably, and that there are many tumors with weaker association between MR and RT. In order to systematically investigate this phenomenon, we divided the genome into four equal bins of RT (limiting the analysis to genomic regions with constitutive RT, see **Methods**), and checked the mutational load in these RT regions for each tumor. Then, we clustered the 2787 WGS tumors into two clusters according to the MR in each bin (**Methods)**. Cluster 1 contains 1042 samples in which the association between MR and RT was weak, while cluster 2 contains the remaining 1745 samples in which the association is stronger (Figure 1 A-B). For simplicity of further analyses, we defined the RT-MRa (“RT-MR association”) metric as the log2 of the ratio between the MR in the late and the early bins 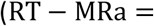 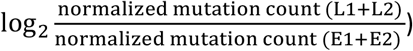. This RT-MRa metric ranges from −1 to 2.5, and most of the samples with RT-MRa score <0.8 were grouped to cluster 1 (Figure 1A (the rightmost column) and 1C).

Next, we examined the possibility of an association of the RT-MRa metric with tumor type (Figure 1D). There are cancer types (i.e., different projects) that contain almost only tumors with weak RT-MRa (cluster 1), such as Kidney Renal Papillary Cell Carcinoma (KIRP) and Breast Cancer (BRCA), while others are heavily biased toward strong RT-MRa (cluster 2), such as Esophageal Adenocarcinoma (ESAD) and Lung Squamous Cell Carcinoma (LUSC). Furthermore, projects of similar cancer types showed similar behavior in this tendency (Figure 1D – different projects of similar tumor types were given the same color).

### Mutational signatures association with the RT-MRa metric

In order to explore the factors that influence RT-MRa, we next analyzed the contribution of mutation signatures. The mutational signatures found in each tumor reflect the mutational processes that the tumor underwent. We have shown that mutational signatures differ in their association with RT [4], and thus they may explain RT-MR association (RT-MRa). We would predict, therefore, that a tumor that mostly underwent mutational processes that are ERR-biased will end up with relatively more mutations in ERR, resulting in a weaker RT-MRa metric as seen in cluster 1.

To explore this relation between mutational processes and RT, we checked the correlation between the RT-MRa metric and the relative contribution of different mutational signatures. We found mutational signatures that display positive correlation with the RT-MRa metric (i.e., samples with larger contribution of these signatures have stronger RT-MRa or belong to cluster 2), and mutational signatures that display negative correlation (i.e., these samples have weaker RT-MRa or belong to cluster 1) (Figure 2A). In some cases, the reason is clear: signatures that are associated with ERR (such as SBS2 and SBS13 [18]) are associated with cluster 1 tumors, whereas signatures associated with LRR (such as SBS17a&b and SBS7a [18]) are associated with cluster 2 tumors. In addition, it is known that defects in DNA repair pathways can cause a different association between RT and MR [5]. Indeed, signatures associated with defects in DNA repair mechanisms (APOBEC, base excision repair, mismatch repair and homologous recombination; Figure 2A - the bold SBS) display negative correlation with RT-MRa, (i.e., contribute more to cluster 1 tumors). Similar association with signatures associated with DNA repair defects was found using the Kruskal-Wallis rank test (see **Methods** and Supplementary Figure S1).

Next, we analyzed the association of the RT-MRa metric with mutational signatures for each project separately. This type of analysis allowed us to identify signatures that are cancer-type specific and thus are less prominent in a pan-cancer analysis. This analysis revealed many signatures that are either positively or negatively associated with the RT-MRa metric in the specific projects (Figure 2B). As expected, signatures with a low mean correlation coefficient value are associated with cluster 1, whereas signatures with a high correlation are associated with cluster 2, which is graphically captured in Figure 2C left panel. The association between RT-MRa and mutational signatures is explained by the RT basis of each signature [4]. Indeed, plotting the average correlation coefficient as a function of the RT bias (delta ERR-LRR) of each signature, demonstrates this association (Figure 2C right panel).

After examining each signature separately, we checked whether combinations of signatures contribute to the RT-MRa. To this end, we represented each tumor by a vector containing the relative contribution of each signature, and used principal component analysis (PCA) followed by K-means clustering to define subgroups of tumors within each project (see **Methods**). Our approach revealed that there are distinct subgroups in many projects, and we examined whether these subgroups are associated with the RT-MRa metric using Wilcoxon rank sum test. In many projects, such as COAD-US, a strong association was found (Figures 2D, Supplementary Figure S3 and Supplementary Table S1). This finding illustrates the importance and impact of the contribution of signatures to the association between RT and mutation rates.

Taken together, mutational processes appear to have a strong association with the RT-MRa in many tumors, yet in many cancer types we found variation in RT-MRa that cannot be explained by signatures (Supplementary Figure S3 and Table S1).

### Identification of pathways significantly mutated in tumors with weak RT-MR association

Next, we explored the possibility that the differences in the RT-MRa scores stem from non-functional components of DNA replication, repair and chromatin structure. To this end, we looked at the abundance of deleterious mutations in tumors with weak and strong RT-MRa (clusters 1 and 2 respectively). Since deleterious mutations are rare, we decided to perform this analysis on a pathway level, meaning that instead of asking whether a mutation in a particular gene is enriched in cluster 1, we asked if mutations in any gene belonging to a particular pathway are enriched in cluster 1. We focused on a few candidate pathways that we hypothesized that they might be associated with changing RT-MRa. These pathways included DNA repair, DNA replication and chromatin organization, and as a control we chose others, non-replication related pathways, as well as random sets of genes. We counted the number of deleterious mutations in each gene in those groups separately for cluster 1 and cluster 2 tumors and calculated the enrichment in cluster 1 using a binomial test (see **Methods**). We found that the DNA repair and chromatin organization pathways were highly enriched for deleterious mutations in cluster 1 samples, whereas no such enrichment was observed in any of the controls (Figure 3A), suggesting that, as expected, those pathways are involved in the modulation of the association between mutation rates and RT. This conclusion was independent of the definition of the deleterious mutations since similar results were observed using different definitions of effective mutations (Supplementary Figure S4). Sub pathways analysis (see **Methods**) did not reveal any informative differences between distinct DNA repair pathways, chromatin modifiers and stages in the DNA replication process (Supplementary Figure S5). The ten most enriched genes in each of the three enriched pathways are shown (Figure 3B and Supplementary Table S2).

### Identification of differentially expressed genes in tumors with weak RT-MR association

In parallel to mutations pathway analyses, we performed differential expression analysis to identify correlation of gene expression and pathways with the RT-MRa metric. This analysis was performed in the 803 samples for which RNA-seq data exists in addition to the WGS mutation information in ICGC [17]. We analyzed each project separately, since different tissues differ in their expression profiles. As we have shown (Figure 1D), the RT-MRa metric varies in most projects and thus can be used to define within each project a set of samples with stronger and weaker association between RT and MR. To this end, we divided the samples in each project into three equally sized groups according to the RT-MRa values and used DESeq2 to identify differential expressed genes between the strong RT-MRa and weak RT-MRa groups (Figure 4A and Supplementary Figure S6). We found numerous genes in most projects that passed the FDR<0.1 criterion. In order to control for inflated FDR in DEseq2 analyses [19], we calculated an experimental FDR by randomizing the tumors in each project, performed differential expression analysis and counted the number of genes that passed the FDR<0.1 threshold in the randomized data. Only projects in which the number of differentially expressed genes in the randomized data that passed the threshold was less than 10% of the number of genes identified by DESeq2 in the original data were considered valid and were kept for further analysis. Following this analysis, we were left with 10 projects of interest. 9 of these had genes expressed higher in the weak RT-MRa group, and 6 had genes expressed higher in the strong RT-MRa group (5 projects contained genes in both groups). We performed GO annotation analysis on these genes using the Metascape tool [20], and identified several categories enriched in the weak RT-MRa group. Interestingly, very similar GO terms were enriched in multiple projects (Figure 4B, Supplementary Tables S3 and S4), suggesting common mechanisms, despite the wide range of tissues and phenotypes. Surprisingly, the enriched reoccurring categories were associated with communications between cells and with the immune system, suggesting that tumors with weak RT-MRa contain a higher degree of infiltration of immune cells (which most probably cause the enrichment of immunological categories). These results were confirmed using a more stringent differential expression identification approach (based on a Wilcoxon test (following [19])) (Supplementary Figure S7, Supplementary Tables S5 and S6). These results suggest that the change in the mutation distribution is not a cell autonomous phenomenon but is rather associated with other cells in the vicinity of the tumor.

Next, we looked at the group of genes that were enriched in several projects. Counting the number of projects each gene was enriched in revealed a large number of genes enriched in multiple projects (Figure 4C). The 400 differential genes that are enriched in at least 3 projects appear mainly in 4 projects – Colon Adenocarcinoma (COAD-US), Head and Neck Squamous Cell Carcinoma (HNSC-US), Head and Neck Thyroid Carcinoma (THCA-US) and Pancreatic Cancer (PACA-CA), (Figure 4D) and are enriched for immunity-associated processes (Figure 4E). Overall, we found that there are several projects where immunity-associated processes seemed to be involved in modulating the association between RT and MR. Samples with higher expression of genes associated with the immune system tend to have weaker associations between RT and MR (regardless of mutation load, see **Discussion**), supporting previous findings regarding the associations between immune processes and replication stress [15, 16].

By contrast, analyzing genes with higher expression in samples with strong RT-MRa did not reveal any association with cell-cell communication or immunology (Supplementary Figure S8, Supplementary Tables S7 and S8).

## Discussion

Replication timing is strongly associated with mutation rates, with more mutations occurring at LRR (reviewed in [2]). Previous studies suggested that this association stems from more efficient DNA repair in the early replicating, more accessible parts of the genome. Indeed, defects in either mismatch repair (MMR) or global genome nucleotide excision repair (GG-NER) mechanisms abolish the RT-MRa [5, 6]. Here we expanded this approach and systematically analyzed the association between RT and MR using 2,787 whole-genome sequenced (WGS) tumors (PCAWG; [17]). We found that there are high levels of variability in RT-MRa between tumors. In approximately 30% of the tumors the association is weak, with almost the same number of mutations in early and late replicating regions. We grouped the samples according to the degree of their RT-MRa in order to identify the molecular processes that modulate the association.

We have previously shown that different mutational processes have distinct associations with RT [4]. Thus, we expected that the combination of signatures characterizing each tumor sample would affect the RT-MRa. Indeed, tumors that underwent mutational processes more abundant in ERR show weaker RT-MRa (**Figure 2**), reflecting the fact that they have a lot of mutations in ERR. This simple explanation for the variability in RT-MRa explains much of the difference in the RT-MRa score between different tumor types (**Figures 1D and 2C**). We also found that mutational signatures related to DNA repair have a greater contribution in tumors with the weaker association, supporting previous observations that defects in DNA repair pathways can affect the association between RT and MR [5, 6]. Overall, clustering the samples based on the mutational signatures results in subgroups that differ in their RT-MRa metric (**Figure 2D**). Yet, in many cases the signatures are not associated with RT-MRa, suggesting that other processes are modulating the genome-wide distribution of mutations.

Next, we were able to show that deleterious mutations in genes of the DNA replication, DNA repair and chromatin organization pathways are enriched for tumor with weaker RT-MRa. Moreover, this enrichment is not limited to the MMR and GG-NER pathways but extends to additional types of DNA repair mechanisms (**Supplementary Figure S5**), as well as replication and chromatin structure pathways, suggesting that multiple cellular processes contribute to the uneven distribution of mutations.

Differential expression analyses revealed that genes highly expressed in the weak RT-MRa samples were enriched in cellular processes involved with interactions between cells and especially with the immune system, and not with the expected cellular processes of DNA replication and repair. Due to the unexpected nature of these results, we took additional measures of precautions. First, we ruled out the possibility of artefacts in the deferential expression analysis by calculating an experimental FDR (based on randomization of the group assignments) and by using a different, more conservative differential expression algorithm (based on a non-parametric test). Secondly, we considered only GO terms that were enriched in several projects (**Figure 4b**). Finally, we found the same pathways enriched for genes that are differentially expressed in several projects (**Figure 4 c-e**). Finding immune genes highly expressed in samples with weak RT-MRa suggests that these samples have higher degrees of immune cells infiltration. This explanation cannot be valid for the Acute Myeloid Leukemia project (LAML-US) since it is not a solid tumor.

Our findings of a correlation between immune cell infiltration with weak RT-MRa can be explained by two opposite explanations. It is possible that cells with weak RT-MRa recruit the immune system with higher efficiency than those with strong RT-MRa; alternatively, it may be that cell-cell interactions, and particularly interactions of the cancer cells with immune cells, affect the genome-wide distribution of mutations. Indeed, tumors with high mutation loads tend to be more immunogenic [21], yet this cannot be the explanation for the enrichment of immune genes in the weak RT-MRa samples, since the mutation load in all those samples (besides in COAD-US) were lower or equal than in the strong RT-MRa group (**Supplementary Figure S9**). Thus, either the non-conventional distribution of mutations recruits the immune system, or the involvement of the immune system somehow modulates mutation distribution.

Both explanations are consistent with recent findings regarding the association of the immune system with replication stress [7, 12–14]. Accumulating evidence indicates that replication stress–inducing agents such as topoisomerase inhibitors and cells deficient in replication stress response genes induce the expression of type I IFNs and proinflammatory cytokines [15, 22–24]. On the other hand, inflammation causes replication stress, by the influence of ISG15 [16]. The fact that there is enrichment of immune genes in the weak RT-MRa samples, raises the question of why is it confined to those samples. This can be explained by the assumption that collision between replication and transcription machineries is the main cause of the immune-related replication stress [25, 26]. Such collisions are expected to be found especially in early replicating regions, due to the prevalence of highly expressed genes in these areas [27–30]. This, in turn, leads to weak RT-MRa [31]. Indeed, a higher percentage of mutations fell within genes in the samples with weak RT-MRa, in most (6/9) of the relevant projects (**Supplementary Figure S10**).

An association between replication stress and inflammation (by IFI16 / STING pathway) has been shown in hidradenitis suppurativa (HS) patients [32]. Yet in the context of cancer, this was demonstrated only in tissue culture systems [7, 15, 16, 22, 24]. To the best of our knowledge, this is the first demonstration of the association between mutation distribution and the immune system in patients’ tumor samples. Further research is needed in order to confirm and to understand the molecular mechanisms underlying this intriguing finding.

## Methods

### Data sources

We downloaded somatic mutation calls (VCF files) from the PCAWG consortium release of 2,787 whole-cancer genomes across 38 tumor types [17]. The data consists of two sources—The International Cancer Genome Consortium (ICGC; 1902 samples) and The Cancer Genome Atlas (TCGA; 885 samples). Each source utilized its standard variant call pipeline (Consensus calls for ICGC, and the Broad Institute variant calling pipeline for TCGA). The somatic mutation profile of the two consortiums were very similar both in terms of 96 trinucleotide context, and in terms of mutational signatures [4]. Accordingly, we combined the mutation calls data for all analyses.

In addition, we downloaded expression data available for 803 of the WGS samples in ICGC. We only included data from primary tumors for the differential expression analysis.

### Cluster division

To minimize the effect of variation in RT between cell types, we only used the constitutive RT regions for our analyses, which constitute approximately 40% of the human genome that have the same RT in 26 tissues examined [33] and are also similar in cancer [4]. Constitutive RT regions were divided into 4 equal bins each spanning the same range of RT: earliest, intermediate early, intermediate late, latest (E1, E2, L1, L2 respectively). Among the constitutive RT regions – 498.6 Mb are defined as E1; 227.6 Mb are defined as E2; 238.4 Mb are defined as L1; and 324.6 Mb as L2. For each tumor, the number of mutations in each of the four regions was counted. These counts were normalized for the region’s sizes and the number of mutations per 100Kb were kept. Thus, each tumor is characterized by a vector of length four, which we used for k-means clustering (Spearman rank correlation method) that divided the data into two different clusters. The RT-MRa (“RT-MR association”) metric is 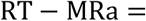 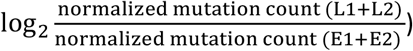

### The association between the signature contribution and clusters tumors

The data of the mutational signatures and their association to RT were taken from Yaacov A, et al. [4]. For each mutational signature we performed correlation test (by the cor.test function in R) with the RT-MRa metric. This was done both for a pool of all tumors (pan-cancer analyses) and for individual projects separately. In the pan-cancer analysis, the test was performed on the tumors in which the signature contribution was above zero. In the cancer-specific analysis, the test was performed on the condition that the mean signature contribution in the project’s tumors was above 5%. The FDR correction was done by the p.adjust (Benjamini-Hochberg Procedure) function in R.

Pan-cancer analyses were also done using ranking statistics. For each mutational signature the tumors were sorted according to the relative contribution of the signature to the overall mutation load. We excluded tumors for which the contribution of the specific signature was below 5%. The association of the rank versus the cluster annotation was assessed using the Kruskal-Wallis rank test. Only statistically significant signatures were shown in Supplementary Figure S1.

For examining the contribution of combinations of signatures to the RT-MRa we represented each tumor sample by the vector of the contribution of each mutational signature to it and used principal components analysis (PCA) to identify groups of similar samples. The resulting groups were further separated by K-means clustering (using the kmeans function in R) to determine the number of clusters by the maximum number of average silhouette widths of the clusters (using the factoextra∷fviz_nbclust R function). We examined the association between our RT-MRa metric and the different groups by Wilcoxon rank-sum test for each pair of clusters. Finally, we computed and plotted the PCA using the stats∷prcomp and ggplot2∷autoplot functions, using the clusters we found.

### Mutational analysis

We focused on four Reactome pathways [34] known to be associated with DNA replication – DNA repair (314 genes), DNA replication (160 genes), chromatin structure (240 genes) and cell cycle (671 genes). As a control we repeated the analysis three times with 500 random chosen genes. In addition, we have chosen some pathways that are less likely to be related with the RT-MRa metric, such as neuronal system, muscle contraction and extracellular matrix organization (containing 419, 196 and 301 genes, respectively). For some of the enriched pathways we also analyzed sub pathways (defined by the Reactome), restricting the analysis to sub pathways containing at least 40 genes.

To focus only on mutations that have a deleterious effect on the protein we used vcf2maf software [35], which provide three different predictions based on Ensembl Variant Effect Predictor (VEP) [36], the Sorting Intolerant from Tolerant (SIFT) algorithm [37] and PolyPhen (Polymorphism Phenotyping) [38].

For each pathway, deleterious mutated genes in each cluster were counted, and then we performed a one-sided binomial test to examine which mutated pathway is enriched in cluster 1. The null hypothesis is that a similar ratio of deleterious mutations between the clusters should be found in all pathways. Thus, we used the ratio in the total deleterious mutations in cluster 1 and 2 as the p in the binomial test. In all cases we corrected for multiple hypotheses testing by FDR.

We performed the same statistical test for all individual genes (Supplementary Table S2). The 10 genes with the lowest P-value are shown in Figure 3B. The mutations in each gene in each cluster was counted and normalized to 100K mutations.

### Differential expression analysis

The differential expression analysis was performed for each cancer project separately. Each project is divided into 3 equal groups according to the RT-MRa score. Then, we used the available expression data from the ICGC to examine which genes are expressed differently between the two extreme groups either by DEseq2 [39] or by Wilcoxon rank-sum test [19]. The Wilcoxon test was performed on a preprocessed count matrix using the edgeR package [40]. The obtained P values of both methods were corrected for multiple hypotheses using FDR. In addition, for the DEseq2 results, we calculated an experimental FDR by randomizing the tumors in each group in each project, re-identifying differentially expressed genes and counting the number of genes that passed the FDR<0.1 criteria in the randomized data.

### GO annotation analysis

For each project we took the genes that were differentially expressed in one of the groups (strong or weak) separately and analyzed it for GO pathways enrichment using Metascape [20]. We limited our analyses to “GO Biological Processes (1148)”. We used the “Heatmap of enriched terms across input gene lists” from the result page. The identification of genes differentially expressed in multiple projects was done using the “Evidence.csv” file (Supplementary Tables S3, S5 and S7). The genes enriched in multiple projects were rerun in Metascape and used the “Heatmap of selected GO”. In addition, we provided the “FINAL_GO.csv” file (Supplementary Tables S4, S6 and S8).

### Statistics

Statistical analyses were performed using R version 4.1.2. For multiple comparisons, P values were corrected by FDR using the p.adjust(method = “BH”) function in R. Plots were generated using ggplot2, ggpubr, and pheatmap R packages

## Acknowledgments

This research was funded by the Israel Academy of Sciences (grant No. 1283/21) and the Binational Science Foundation (grant No. 2019688).

The authors thank Dr. Sheera Adar and May Merav for assistance in data analysis and for critical reading of the manuscript, and to Avraham Greenberg for assistance in editing and writing.

---

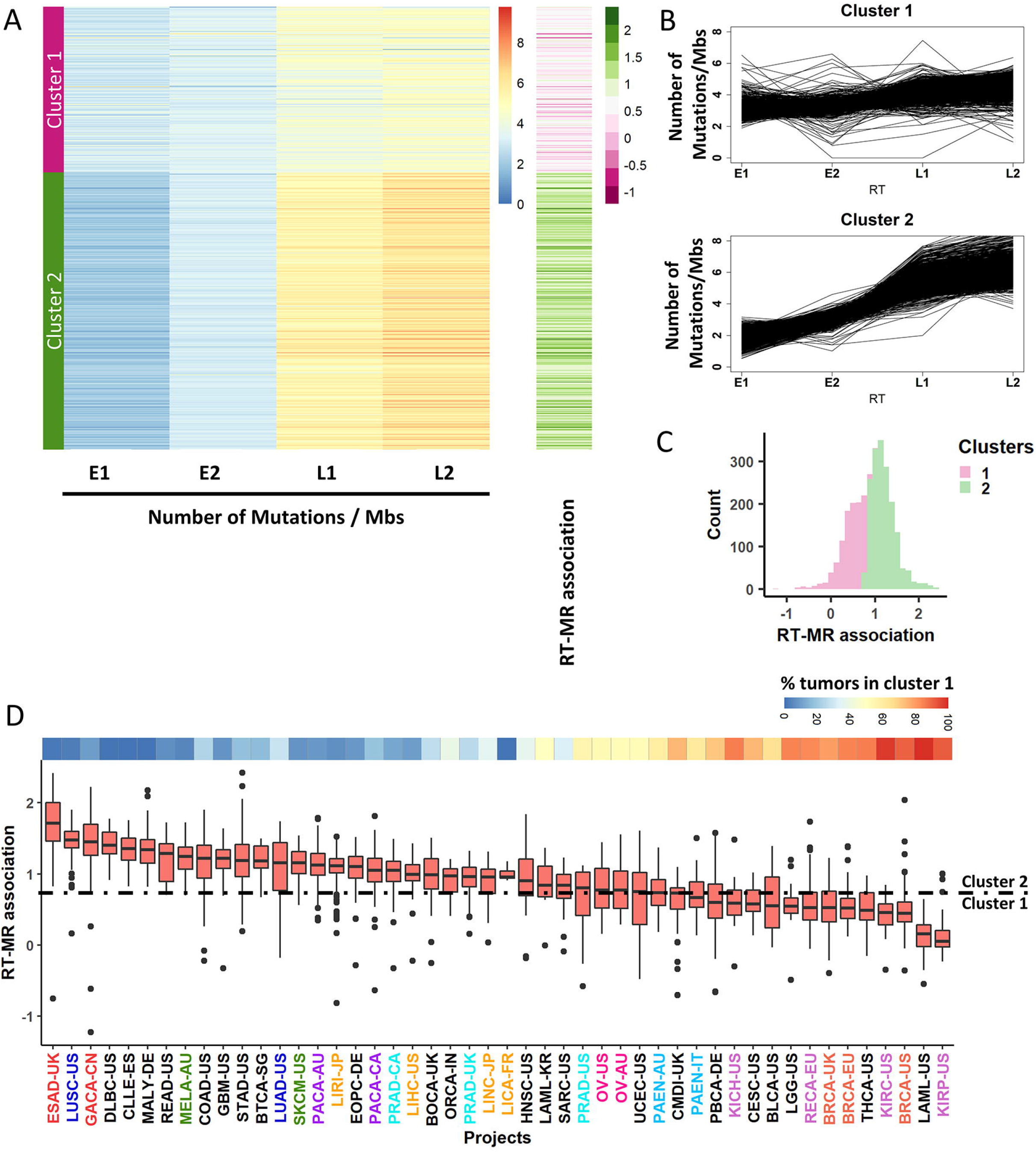

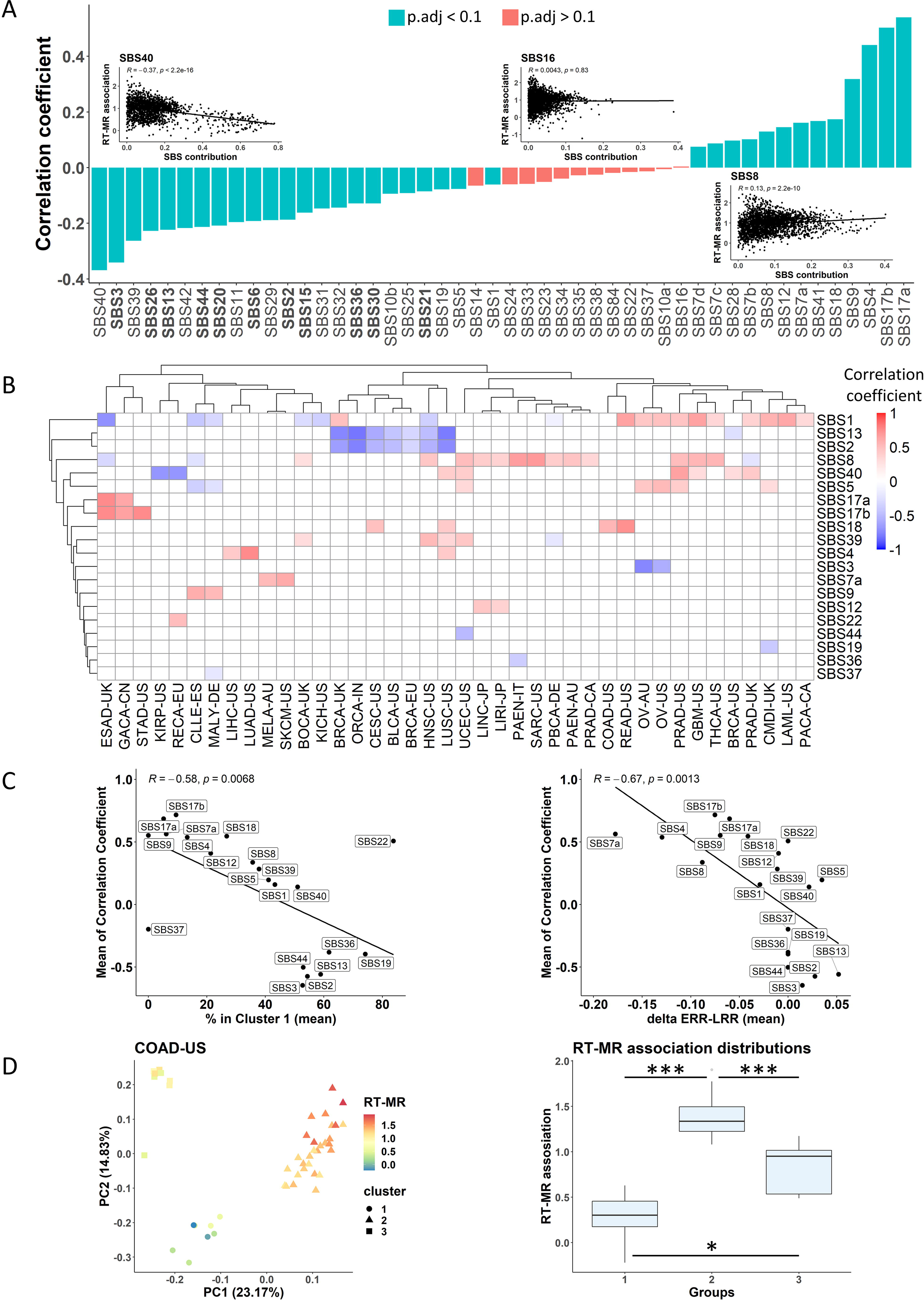

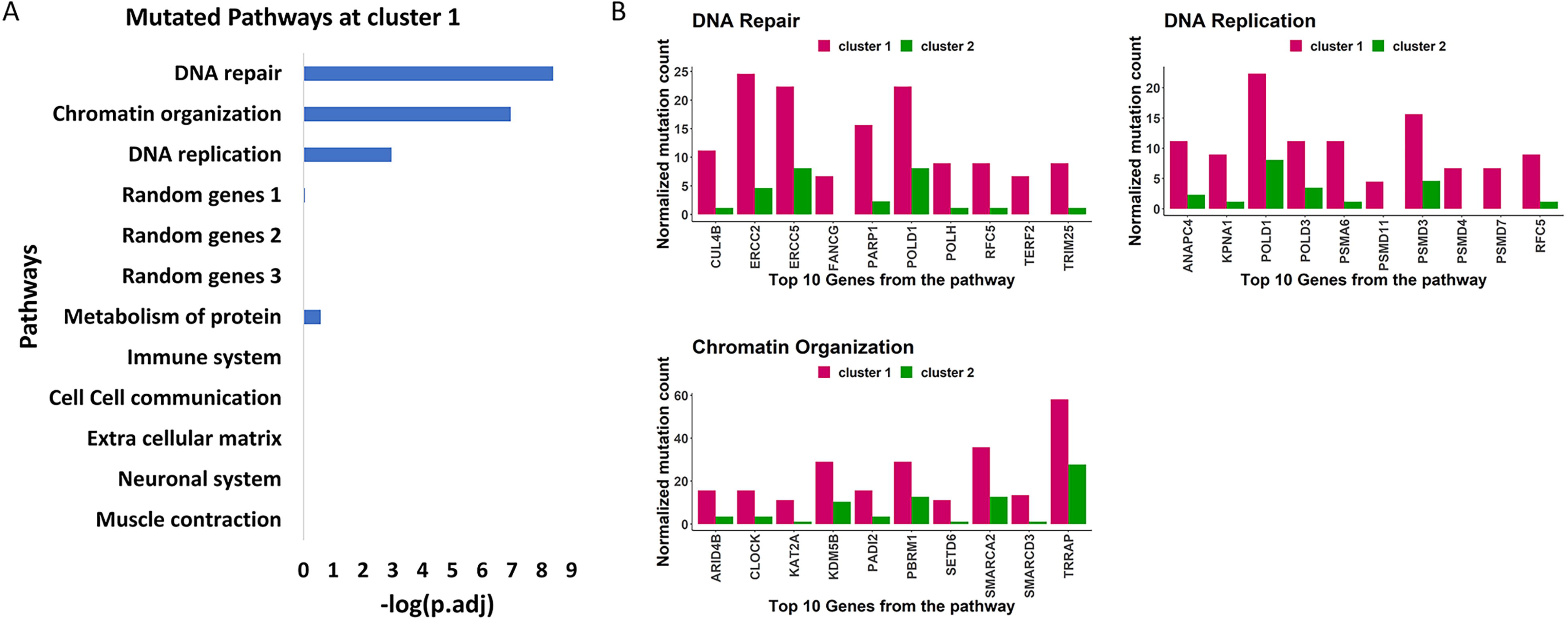

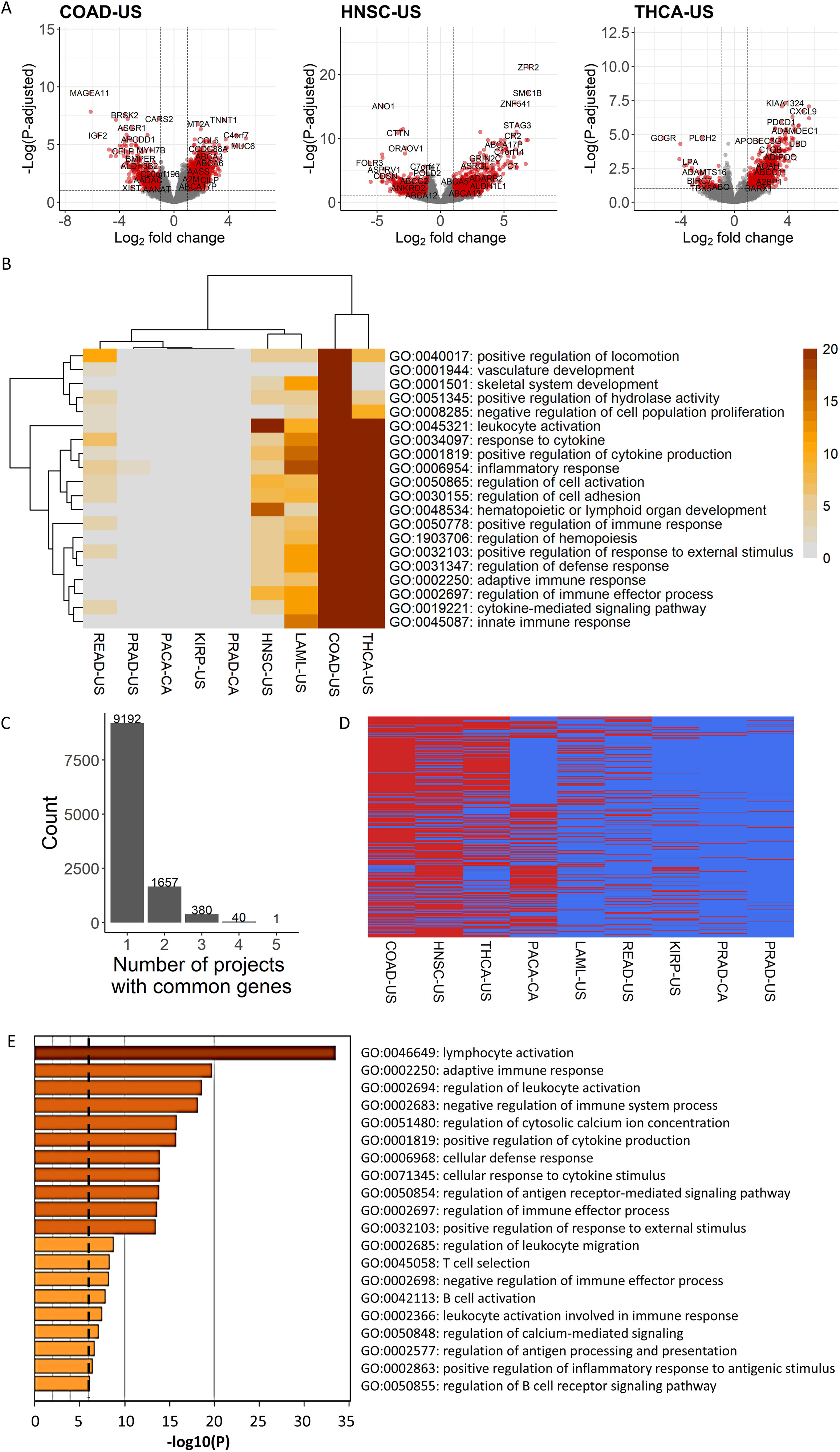

## References

1. Tomasetti, C. and B. Vogelstein, Cancer etiology. Variation in cancer risk among tissues can be explained by the number of stem cell divisions. Science, 2015. 347(6217): p. 78–81.

2. Blumenfeld, B., M. Ben-Zimra, and I. Simon, Perturbations in the Replication Program Contribute to Genomic Instability in Cancer. Int J Mol Sci, 2017. 18(6).

3. Pleasance, E.D., et al., A comprehensive catalogue of somatic mutations from a human cancer genome. Nature, 2010. 463(7278): p. 191–6.

4. Yaacov, A., et al., Cancer Mutational Processes Vary in Their Association with Replication Timing and Chromatin Accessibility. Cancer Res, 2021. 81(24): p. 6106–6116.

5. Supek, F. and B. Lehner, Differential DNA mismatch repair underlies mutation rate variation across the human genome. Nature, 2015. 521(7550): p. 81–4.

6. Zheng, C.L., et al., Transcription restores DNA repair to heterochromatin, determining regional mutation rates in cancer genomes. Cell Rep, 2014. 9(4): p. 1228–34.

7. Bianco, J.N., et al., Overexpression of Claspin and Timeless protects cancer cells from replication stress in a checkpoint-independent manner. Nat Commun, 2019. 10(1): p. 910.

8. Ubhi, T. and G.W. Brown, Exploiting DNA Replication Stress for Cancer Treatment. Cancer Res, 2019. 79(8): p. 1730–1739.

9. Riaz, N., et al., The role of neoantigens in response to immune checkpoint blockade. Int Immunol, 2016. 28(8): p. 411–9.

10. Matsushita, H., et al., Cancer exome analysis reveals a T-cell-dependent mechanism of cancer immunoediting. Nature, 2012. 482(7385): p. 400–4.

11. Thomas, A., et al., Therapeutic targeting of ATR yields durable regressions in small cell lung cancers with high replication stress. Cancer Cell, 2021. 39(4): p. 566–579 e7.

12. Techer, H. and P. Pasero, The Replication Stress Response on a Narrow Path Between Genomic Instability and Inflammation. Front Cell Dev Biol, 2021. 9: p. 702584.

13. Bronner, C.E., et al., Mutation in the DNA mismatch repair gene homologue hMLH1 is associated with hereditary non-polyposis colon cancer. Nature, 1994. 368(6468): p. 258–61.

14. Yu, Q., et al., DNA-damage-induced type I interferon promotes senescence and inhibits stem cell function. Cell Rep, 2015. 11(5): p. 785–797.

15. Coquel, F., et al., SAMHD1 acts at stalled replication forks to prevent interferon induction. Nature, 2018. 557(7703): p. 57–61.

16. Raso, M.C., et al., Interferon-stimulated gene 15 accelerates replication fork progression inducing chromosomal breakage. J Cell Biol, 2020. 219(8).

17. Consortium, I.T.P.-C.A.o.W.G., Pan-cancer analysis of whole genomes. Nature, 2020. 578(7793): p. 82–93.

18. Mas-Ponte, D. and F. Supek, DNA mismatch repair promotes APOBEC3-mediated diffuse hypermutation in human cancers. Nat Genet, 2020. 52(9): p. 958–968.

19. Li, Y., et al., Exaggerated false positives by popular differential expression methods when analyzing human population samples. Genome Biol, 2022. 23(1): p. 79.

20. Zhou, Y., et al., Metascape provides a biologist-oriented resource for the analysis of systems-level datasets. Nat Commun, 2019. 10(1): p. 1523.

21. Wu, Z., et al., Identification of gene expression profiles and immune cell infiltration signatures between low and high tumor mutation burden groups in bladder cancer. Int J Med Sci, 2020. 17(1): p. 89–96.

22. Luthra, P., et al., Topoisomerase II Inhibitors Induce DNA Damage-Dependent Interferon Responses Circumventing Ebola Virus Immune Evasion. mBio, 2017. 8(2).

23. Dunphy, G., et al., Non-canonical Activation of the DNA Sensing Adaptor STING by ATM and IFI16 Mediates NF-kappaB Signaling after Nuclear DNA Damage. Mol Cell, 2018. 71(5): p. 745–760 e5.

24. Shen, J., et al., PARPi Triggers the STING-Dependent Immune Response and Enhances the Therapeutic Efficacy of Immune Checkpoint Blockade Independent of BRCAness. Cancer Res, 2019. 79(2): p. 311–319.

25. Deshpande, A.M. and C.S. Newlon, DNA replication fork pause sites dependent on transcription. Science, 1996. 272(5264): p. 1030–3.

26. Liu, Y., et al., Topoisomerase I prevents transcription-replication conflicts at transcription termination sites. Mol Cell Oncol, 2020. 8(1): p. 1843951.

27. Lalonde, M., et al., Consequences and Resolution of Transcription-Replication Conflicts. Life (Basel), 2021. 11(7).

28. Marchal, C., J. Sima, and D.M. Gilbert, Control of DNA replication timing in the 3D genome. Nat Rev Mol Cell Biol, 2019. 20(12): p. 721–737.

29. Garcia-Muse, T. and A. Aguilera, Transcription-replication conflicts: how they occur and how they are resolved. Nat Rev Mol Cell Biol, 2016. 17(9): p. 553–63.

30. Hamperl, S., et al., Transcription-Replication Conflict Orientation Modulates R-Loop Levels and Activates Distinct DNA Damage Responses. Cell, 2017. 170(4): p. 774–786 e19.

31. Sankar, T.S., et al., The nature of mutations induced by replication-transcription collisions. Nature, 2016. 535(7610): p. 178–81.

32. Orvain, C., et al., Hair follicle stem cell replication stress drives IFI16/STING-dependent inflammation in hidradenitis suppurativa. J Clin Invest, 2020. 130(7): p. 3777–3790.

33. Rivera-Mulia, J.C., et al., Dynamic changes in replication timing and gene expression during lineage specification of human pluripotent stem cells. Genome Res, 2015. 25(8): p. 1091–103.

34. Gillespie, M., et al., The reactome pathway knowledgebase 2022. Nucleic Acids Res, 2022. 50(D1): p. D687–D692.

35. Kandoth, C., mskcc/vcf2maf: vcf2maf. 2020.

36. McLaren, W., et al., The Ensembl Variant Effect Predictor. Genome Biol, 2016. 17(1): p. 122.

37. Sim, N.L., et al., SIFT web server: predicting effects of amino acid substitutions on proteins. Nucleic Acids Res, 2012. 40(Web Server issue): p. W452–7.

38. Adzhubei, I., D.M. Jordan, and S.R. Sunyaev, Predicting functional effect of human missense mutations using PolyPhen-2. Curr Protoc Hum Genet, 2013. **Chapter** 7: p. Unit7 20.

39. Love, M.I., W. Huber, and S. Anders, Moderated estimation of fold change and dispersion for RNA-seq data with DESeq2. Genome Biol, 2014. 15(12): p. 550.

40. Robinson, M.D., D.J. McCarthy, and G.K. Smyth, edgeR: a Bioconductor package for differential expression analysis of digital gene expression data. Bioinformatics, 2010. 26(1): p. 139–40.

