## Supplementary Figures for "Both cell autonomous and non-autonomous processes modulate the association between replication timing and mutation rate"

#### Supplementary Figure S1

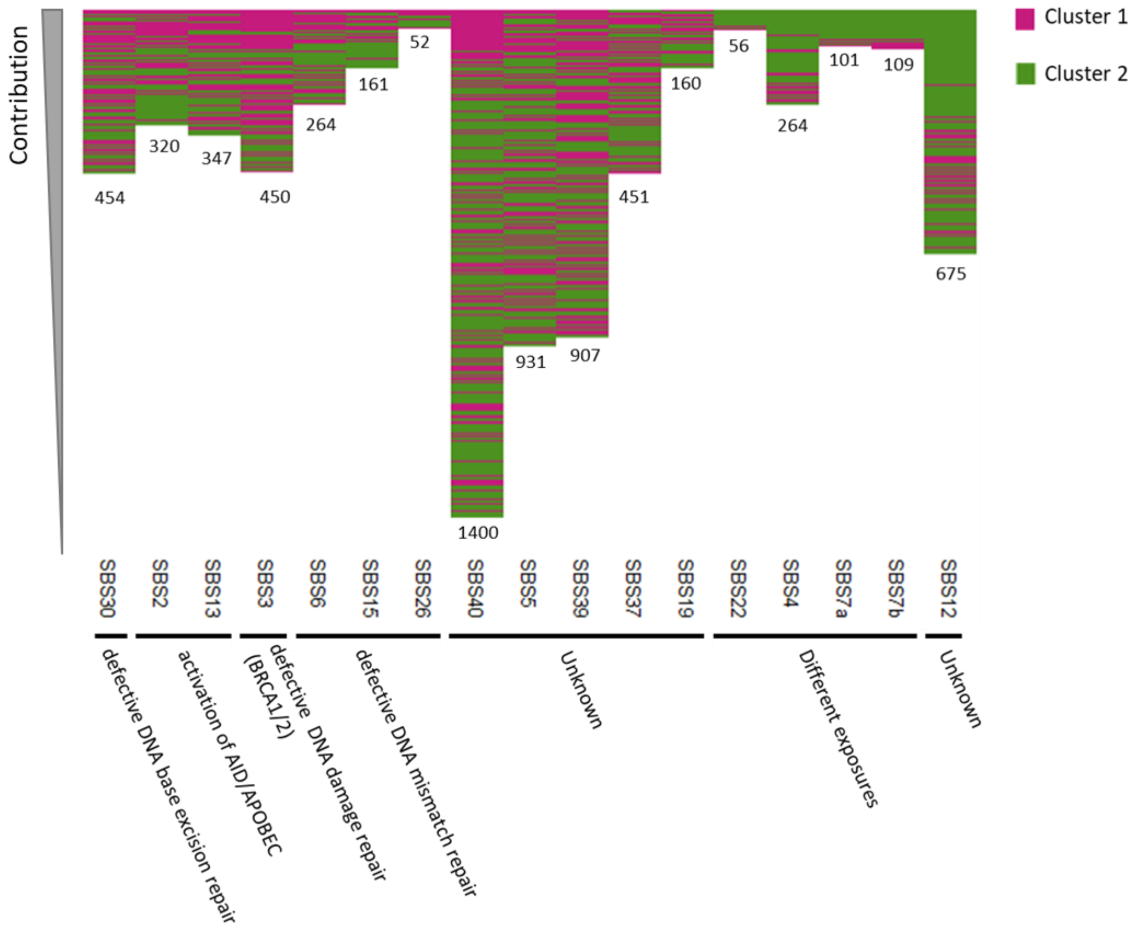


**Supplementary Figure S1. Association between RT-MR metric and Mutational signatures**

For each mutational signature the samples were sorted according to the relative contribution of the signature to the overall mutation load. Only samples in which the signature contribution is above 5% are shown. Samples belonging to cluster 1 are labeled in pink and samples from cluster 2 are green. Signatures with significant association between cluster assignment and contribution are shown. Only statistically significant signatures (adjusted P-val < 0.1; Kruskal-Wallis rank test). As indicated by the colors the left 11 signatures are enriched in cluster 1 samples, whereas the right 5 signatures are enriched in cluster 2 samples. Note that signatures enriched for cluster 1 are mainly associated with defects in DNA repair mechanisms.

### Supplementary Figure S2

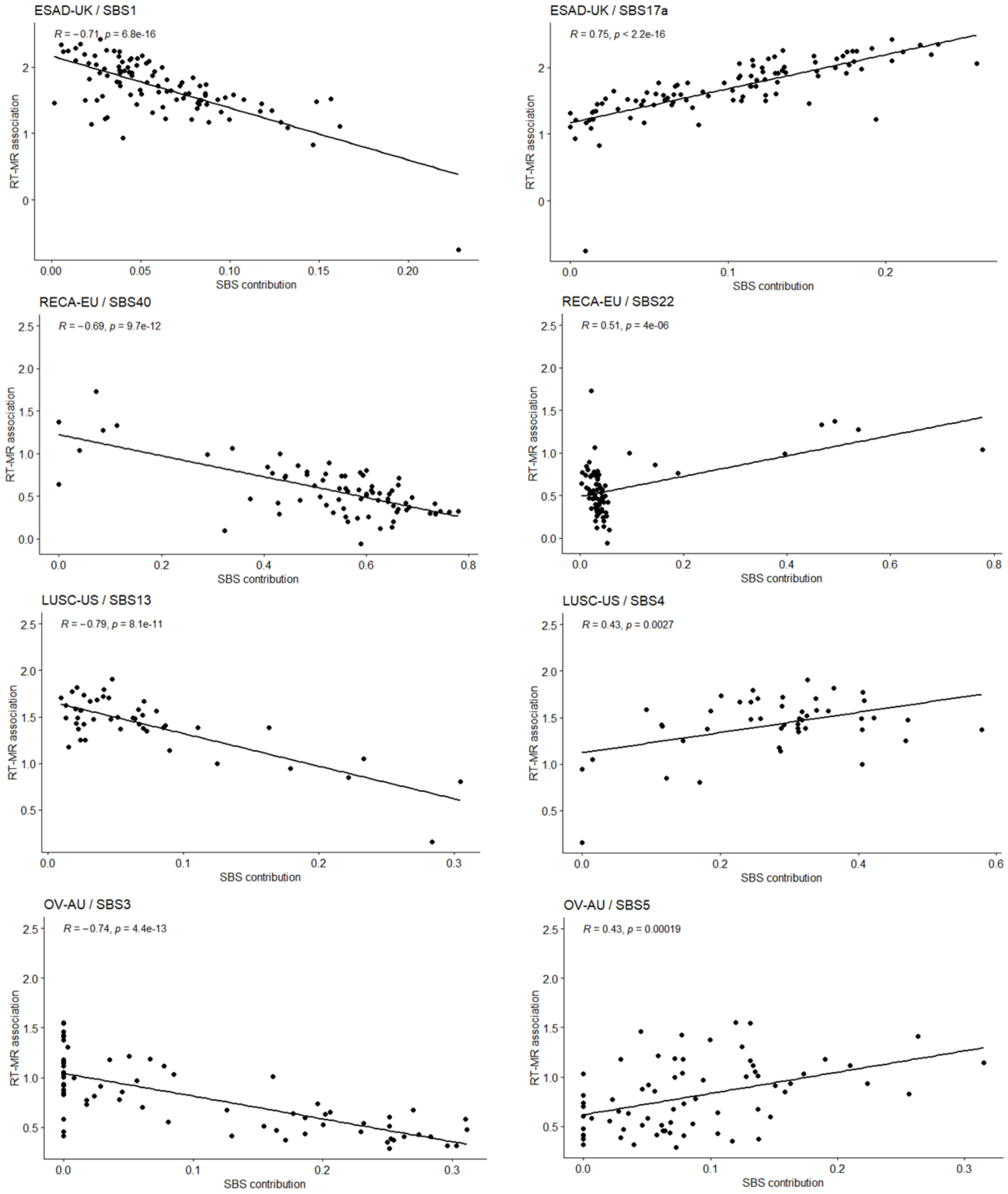


**Supplementary Figure S2. Association between RT-MR metric and Mutational signatures of specific projects**

Scatter-plots showing the association between RT-MR metric and signatures contribution for four projects; ESAD-UK, RECA-EU, LUSC-US and OV-AU. For each project two signatures are displayed – one with positive correlation (mark in red color in Fig. 2B) and one with negative correlation (mark in blue color in Fig. 2B).

### Supplementary Figure S3

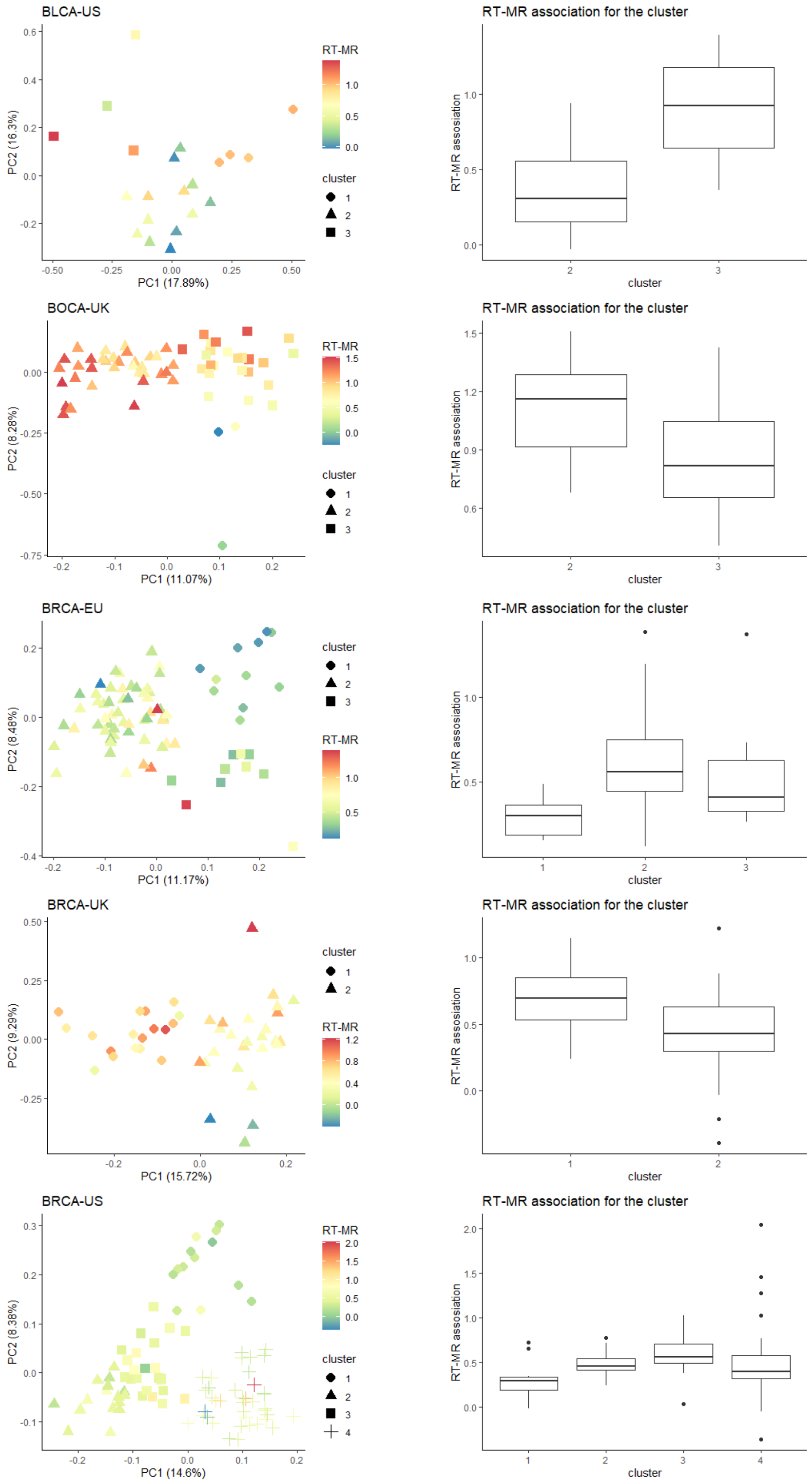

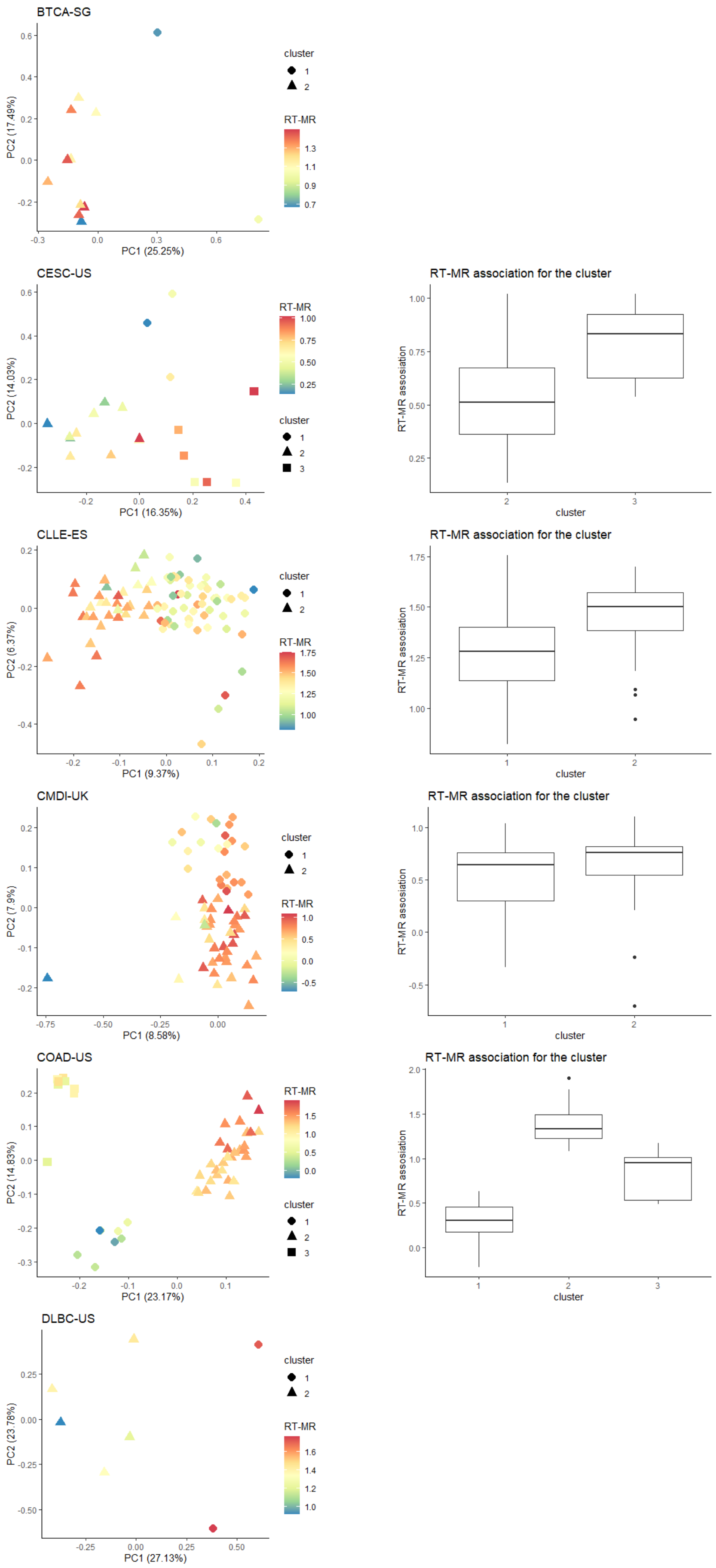

BTCA-SG
PC2 (17.49%)
PC1 (25.25%)
cluster
RT-MR
CESC-US
PC2 (14.03%)
PC1 (16.35%)
RT-MR
cluster
RT-MR association for the cluster
RT-MR assosiation
cluster
CLLE-ES
PC2 (6.37%)
PC1 (9.37%)
cluster
RT-MR
RT-MR association for the cluster
RT-MR assosiation
cluster
CMDI-UK
PC2 (7.9%)
PC1 (8.58%)
cluster
RT-MR
RT-MR association for the cluster
RT-MR assosiation
cluster
COAD-US
PC2 (14.83%)
PC1 (23.17%)
RT-MR
cluster
RT-MR association for the cluster
RT-MR assosiation
cluster
DLBC-US
PC2 (23.78%)
PC1 (27.13%)
cluster
RT-MR

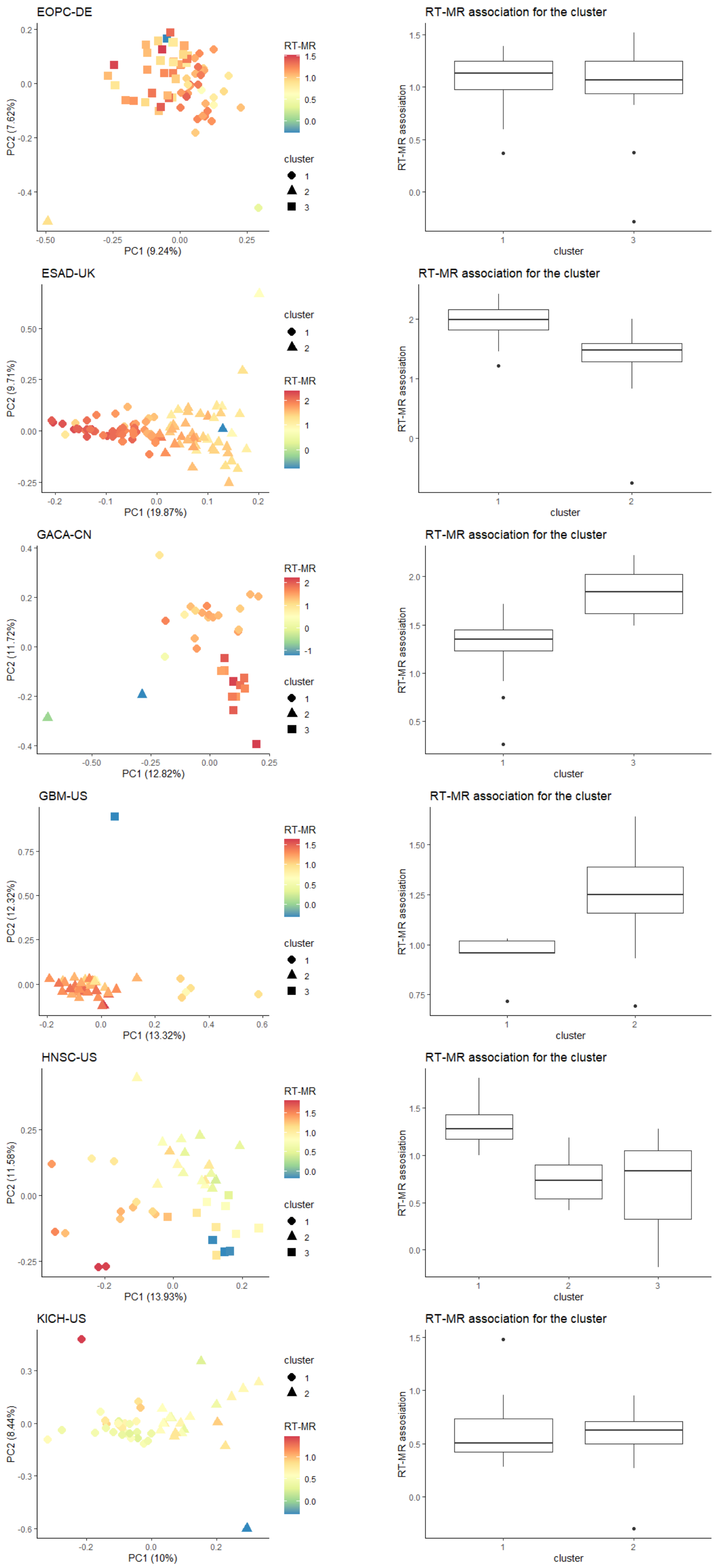
EOPC-DE
PC1 (9.24%)
PC2 (7.62%)
RT-MR
cluster
RT-MR association for the cluster
RT-MR association
cluster
ESAD-UK
PC1 (19.87%)
PC2 (9.71%)
cluster
RT-MR
RT-MR association for the cluster
RT-MR association
cluster
GACA-CN
PC1 (12.82%)
PC2 (11.72%)
RT-MR
cluster
RT-MR association for the cluster
RT-MR association
cluster
GBM-US
PC1 (13.32%)
PC2 (12.32%)
RT-MR
cluster
RT-MR association for the cluster
RT-MR association
cluster
HNSC-US
PC1 (13.93%)
PC2 (11.58%)
RT-MR
cluster
RT-MR association for the cluster
RT-MR association
cluster
KICH-US
PC1 (10%)
PC2 (8.44%)
cluster
RT-MR
RT-MR association for the cluster
RT-MR association
cluster

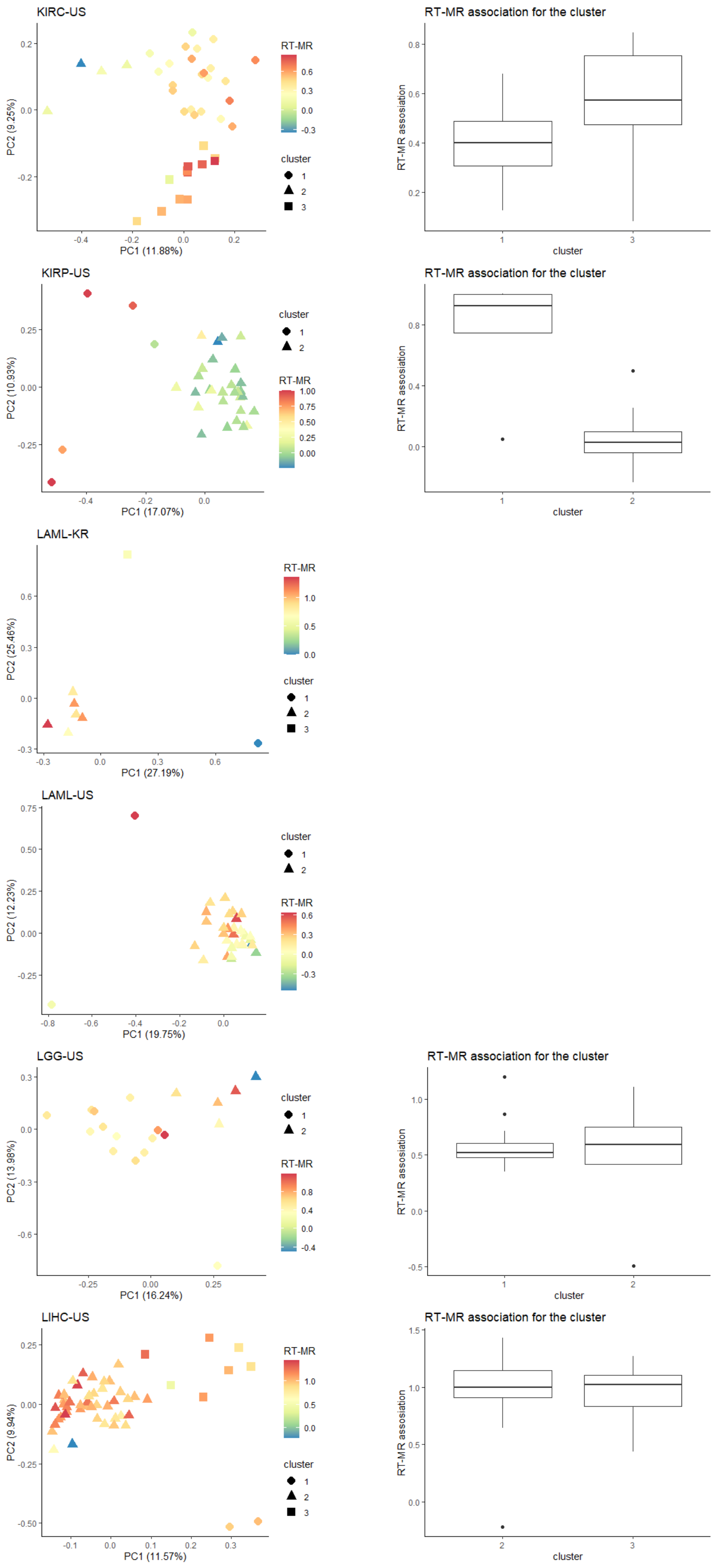

KIRC-US
PC1 (11.88%)
PC2 (9.25%)
RT-MR
cluster
RT-MR association for the cluster
RT-MR association
cluster
KIRP-US
PC1 (17.07%)
PC2 (10.93%)
cluster
RT-MR
RT-MR association for the cluster
RT-MR association
cluster
LAML-KR
PC1 (27.19%)
PC2 (25.46%)
RT-MR
cluster
LAML-US
PC1 (19.75%)
PC2 (12.23%)
cluster
RT-MR
LGG-US
PC1 (16.24%)
PC2 (13.98%)
cluster
RT-MR
RT-MR association for the cluster
RT-MR association
cluster
LIHC-US
PC1 (11.57%)
PC2 (9.94%)
RT-MR
cluster
RT-MR association for the cluster
RT-MR association
cluster

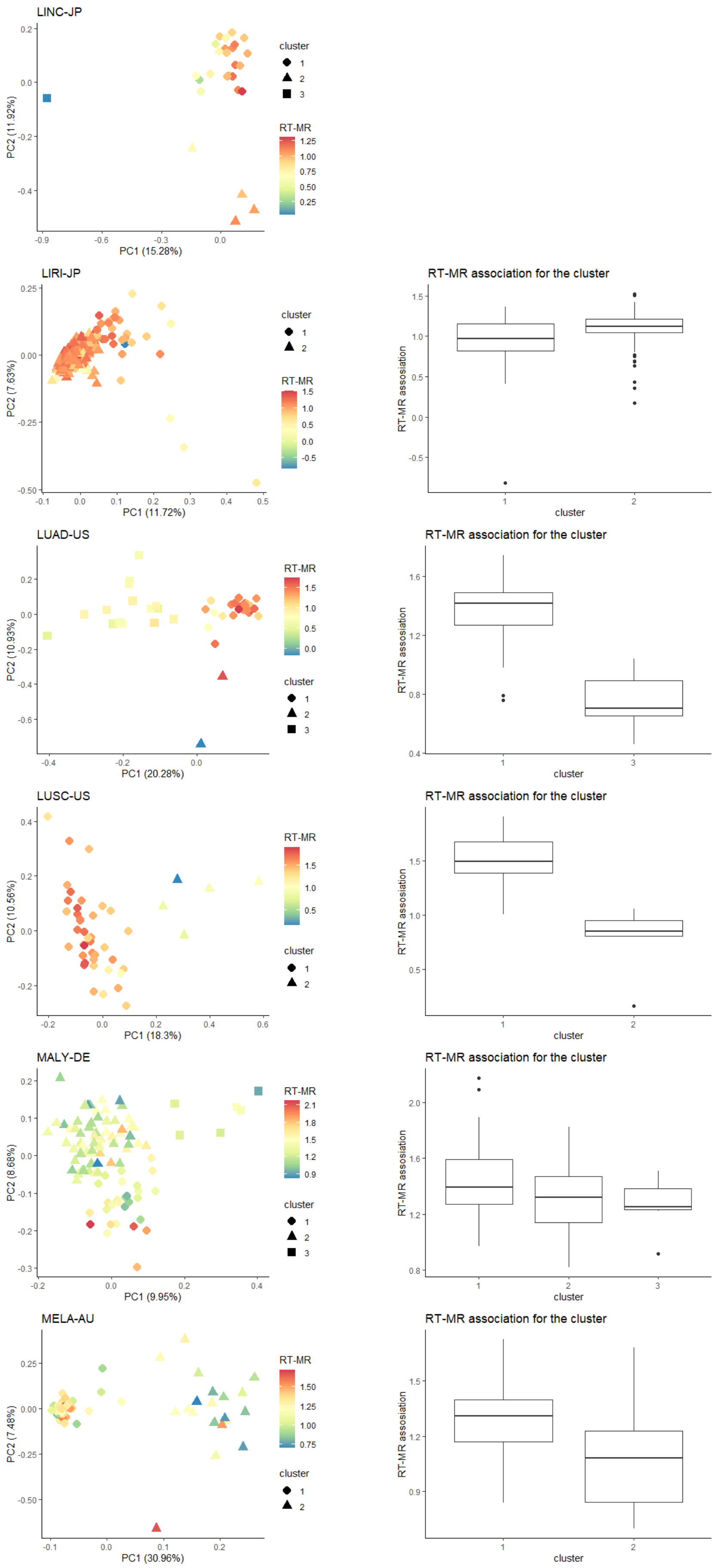
LINC-JP
PC1 (15.28%)
PC2 (11.92%)
cluster
1
2
3
RT-MR
LIRI-JP
PC1 (11.72%)
PC2 (7.63%)
RT-MR association for the cluster
RT-MR assosiation
cluster
LUAD-US
PC1 (20.28%)
PC2 (10.93%)
LUSC-US
PC1 (18.3%)
PC2 (10.56%)
MALY-DE
PC1 (9.95%)
PC2 (8.68%)
MELA-AU
PC1 (30.96%)
PC2 (7.48%)

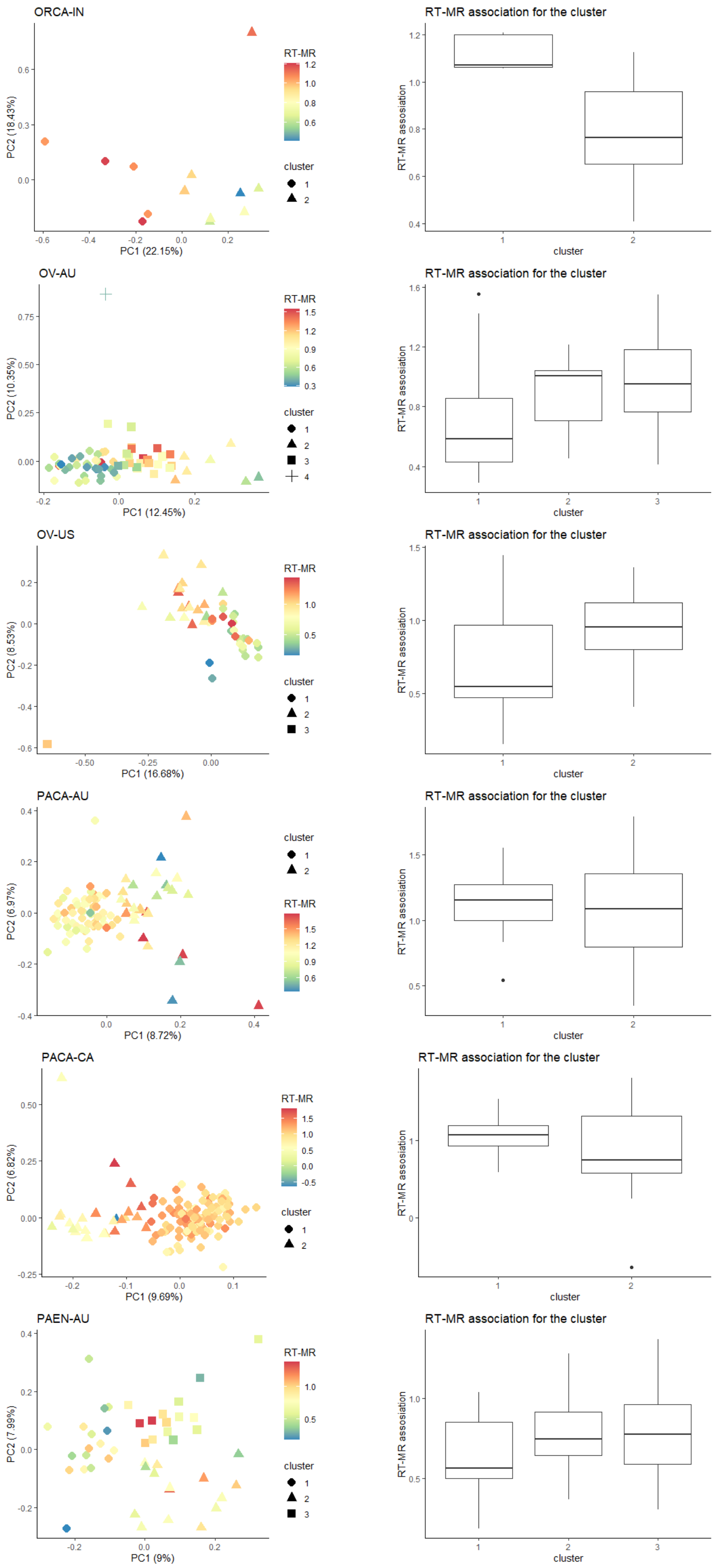

ORCA-IN
PC2 (18.43%)
PC1 (22.15%)
RT-MR
cluster
RT-MR association for the cluster
RT-MR assosiation
cluster
OV-AU
PC2 (10.35%)
PC1 (12.45%)
RT-MR
cluster
RT-MR association for the cluster
RT-MR assosiation
cluster
OV-US
PC2 (8.53%)
PC1 (16.68%)
RT-MR
cluster
RT-MR association for the cluster
RT-MR assosiation
cluster
PACA-AU
PC2 (6.97%)
PC1 (8.72%)
cluster
RT-MR
RT-MR association for the cluster
RT-MR assosiation
cluster
PACA-CA
PC2 (6.82%)
PC1 (9.69%)
RT-MR
cluster
RT-MR association for the cluster
RT-MR assosiation
cluster
PAEN-AU
PC2 (7.99%)
PC1 (9%)
RT-MR
cluster
RT-MR association for the cluster
RT-MR assosiation
cluster

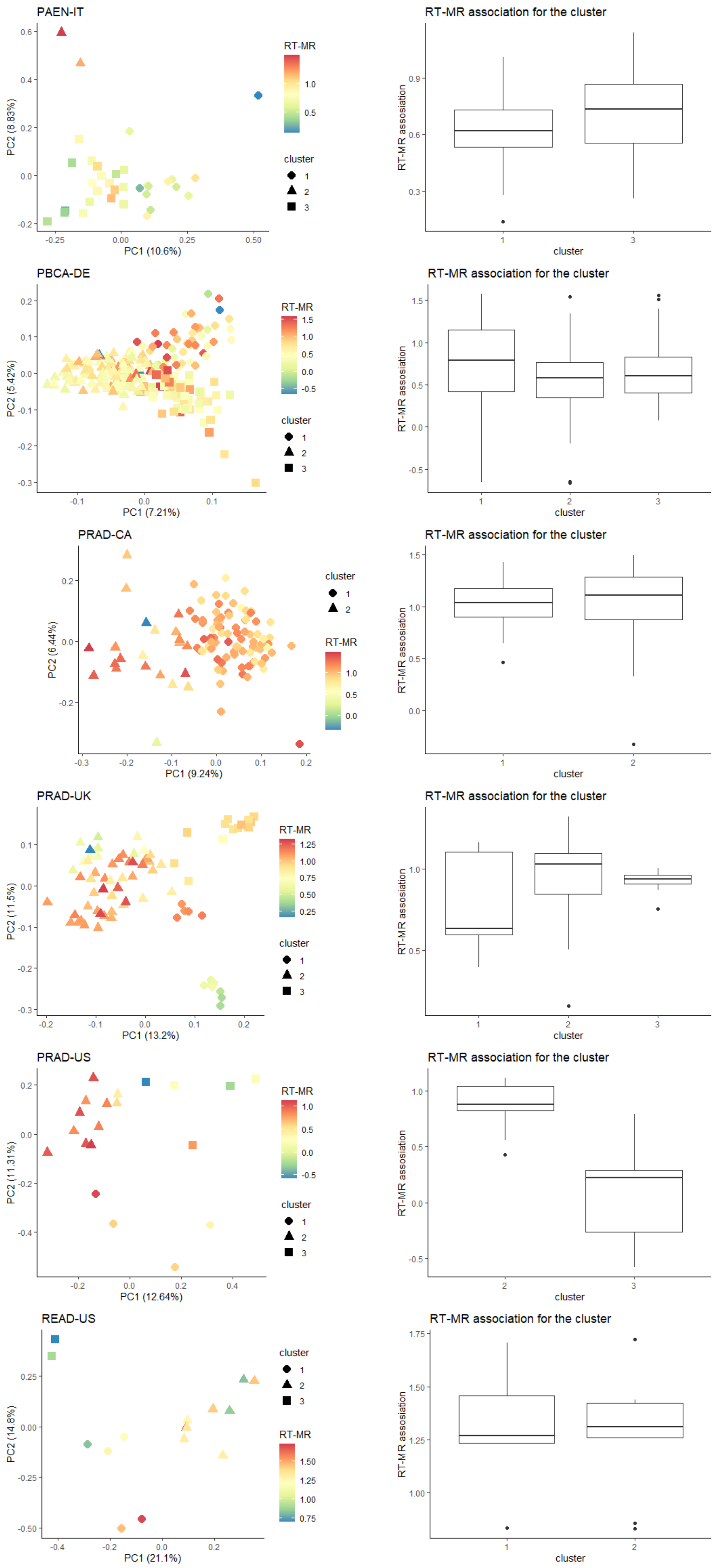
PAEN-IT
PC1 (10.6%)
PC2 (8.83%)
RT-MR
cluster
RT-MR association for the cluster
RT-MR assosiation
cluster
PBCA-DE
PC1 (7.21%)
PC2 (5.42%)
RT-MR
cluster
RT-MR association for the cluster
RT-MR assosiation
cluster
PRAD-CA
PC1 (9.24%)
PC2 (6.44%)
cluster
RT-MR
RT-MR association for the cluster
RT-MR assosiation
cluster
PRAD-UK
PC1 (13.2%)
PC2 (11.5%)
RT-MR
cluster
RT-MR association for the cluster
RT-MR assosiation
cluster
PRAD-US
PC1 (12.64%)
PC2 (11.31%)
RT-MR
cluster
RT-MR association for the cluster
RT-MR assosiation
cluster
READ-US
PC1 (21.1%)
PC2 (14.8%)
cluster
RT-MR
RT-MR association for the cluster
RT-MR assosiation
cluster

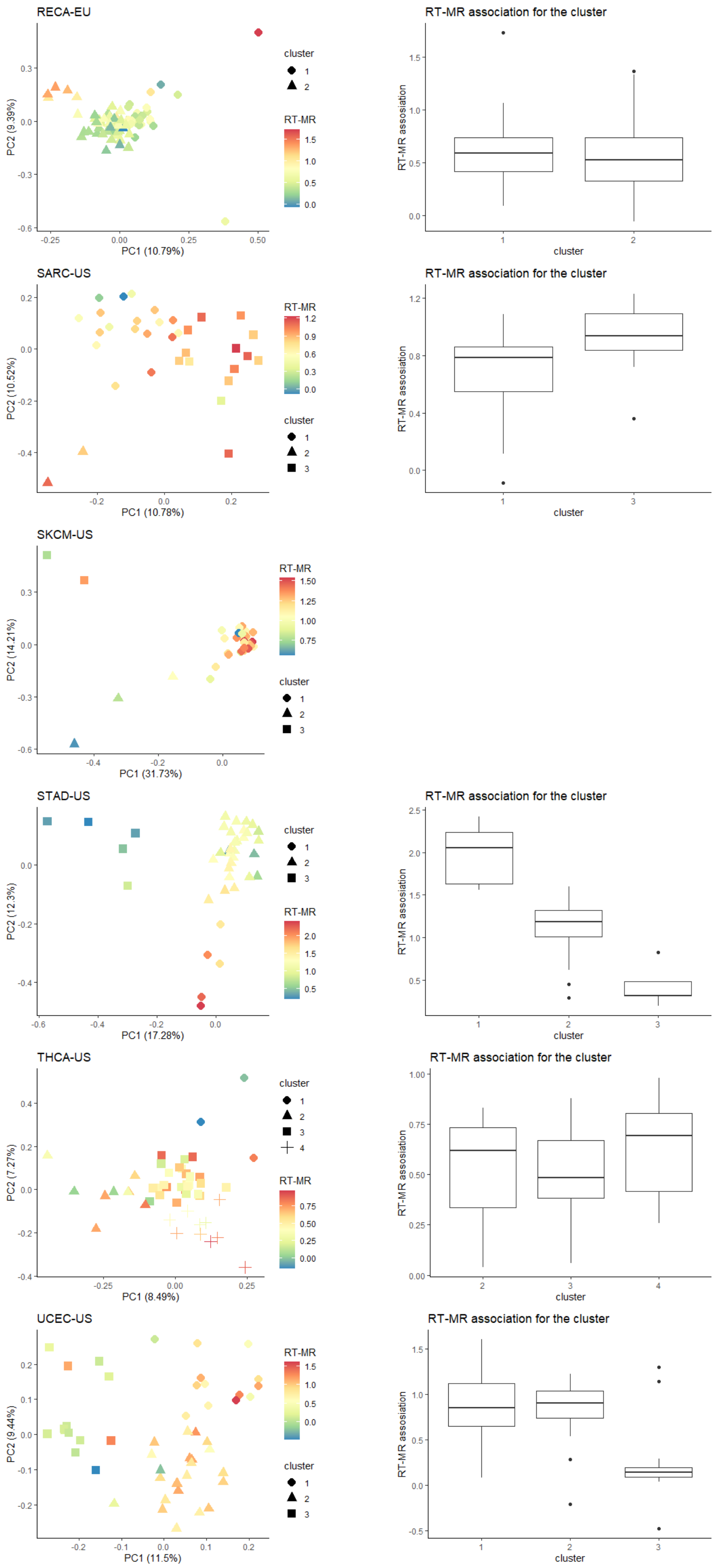


**Supplementary Figure S3. Contribution of combinations of signatures to the RT-MR associations**

For each project shown PCA plots of signature contribution data, colored by RT-MR association score and divided into different shapes by K mean clustering (left panel), and boxplot of the RT-MR association distribution of the different clusters (only for clusters with n>4). All P values derived from FDR-corrected Wilcoxon rank–sum test and shown in **Supplementary Table S1**.

### Supplementary Figure S4

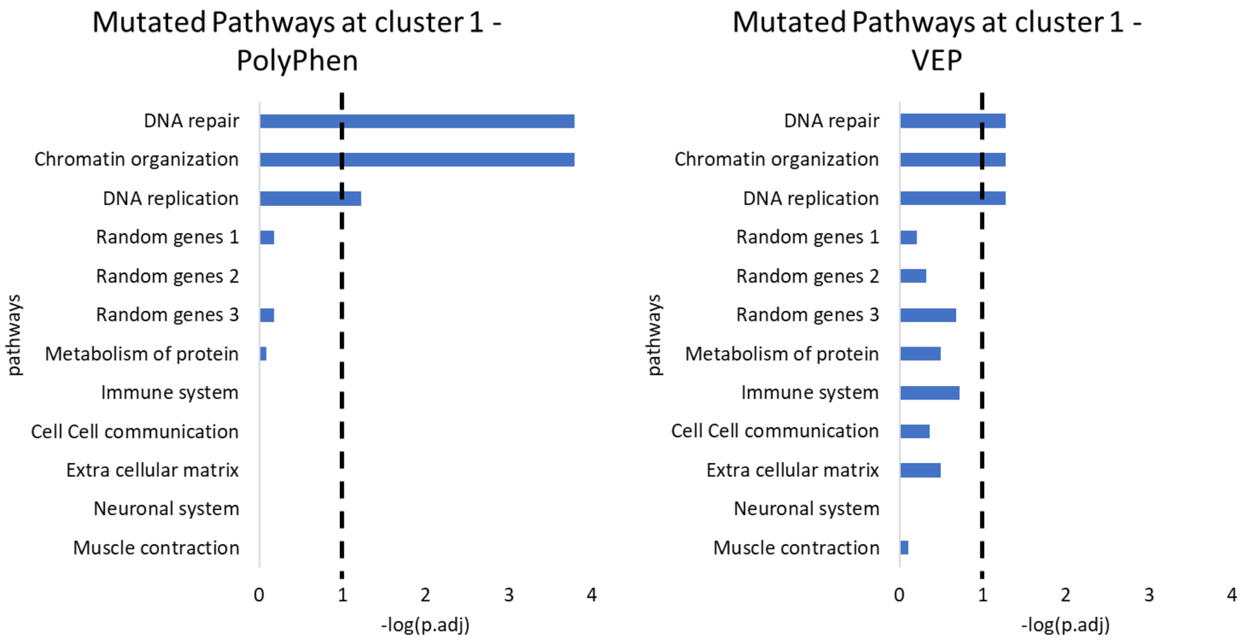


**Supplementary Figure S4. Mutated pathway analysis using different definition of deleterious genes**

The different pathways are tested to examine the enrichment to cluster 1 tumors, using mutation defined as harmful by PolyPhen (left) or by VEP (right). FDR corrected P values (Binomial tests) are shown. The vertical dashed line designates a threshold of adjusted P val<0.1. Note, that in the VEP based dataset the DNA repair category was not significantly enriched in cluster 1 tumors unless mutations in TP53 were removed from the analysis. This was done since TP53 show an opposite trend then the other dana repair genes.

### Supplementary Figure S5

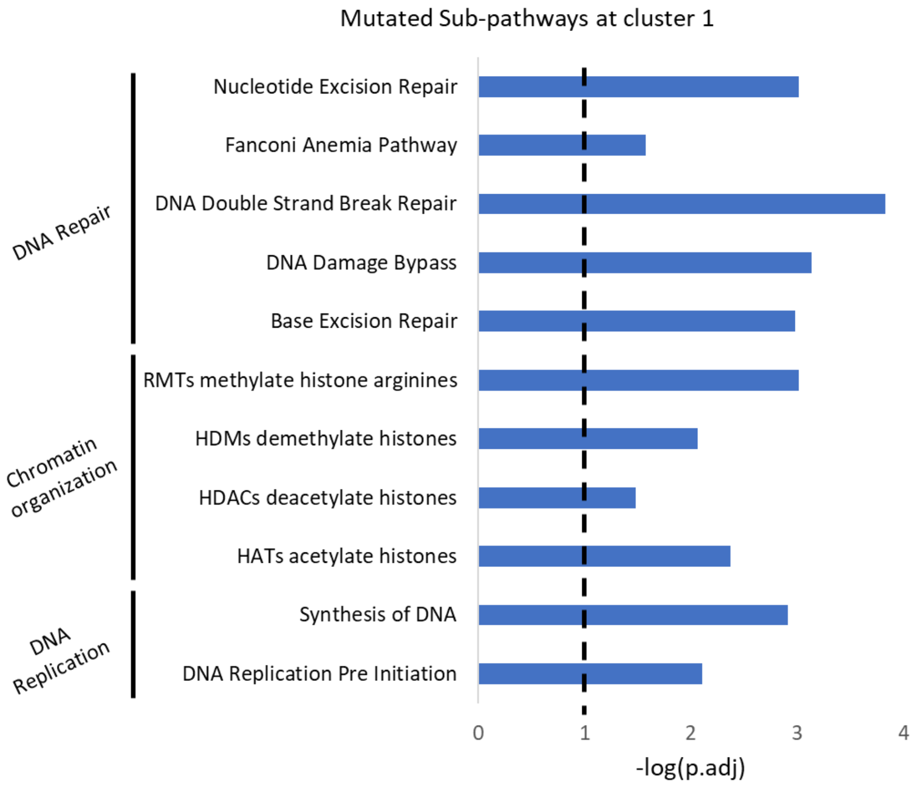


**Supplementary Figure S5. Sub-pathways mutation analysis**

Each of the pathways shown in figure 3A were further divided into sub pathways (using Reactome definition) and the enrichment analyses were repeated using the SIFT definition for deleterious mutations. Only sub-pathways containing more than 50 genes were analyzed. The vertical dashed line designates a threshold of adjusted P-val<0.1. Here also TP53 mutations were removed from the analyses.

### Supplementary Figure S6

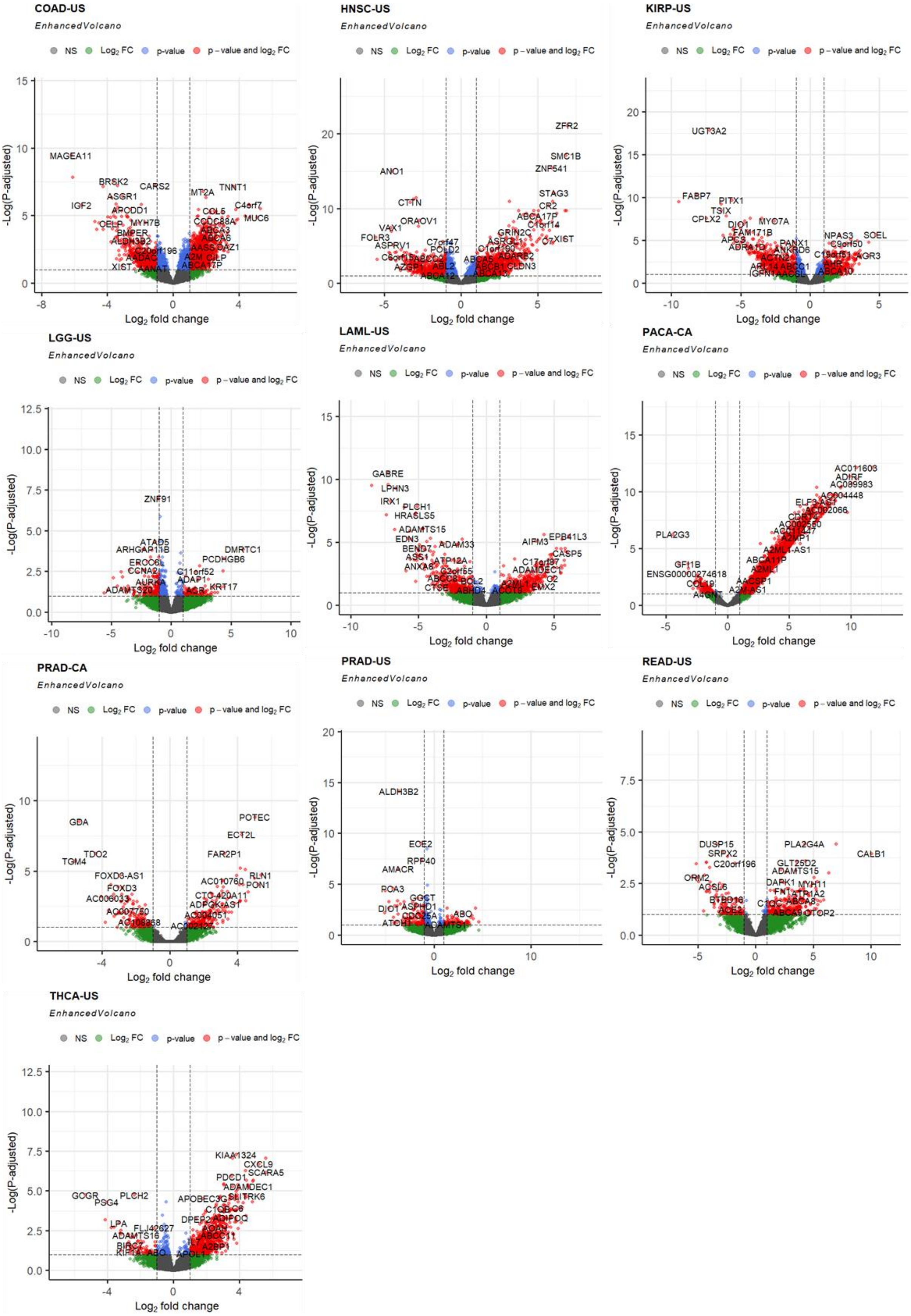

**Supplementary Figure S6. Expression profile analysis**

Volcano-plots of ten significant projects (projects in which the number of differentials expressed genes in the randomized data that passed the threshold was less than 10% of the number of genes identified by DESeq2). The red points are genes that are differentially expressed significantly between the low/high RT-MR groups. These points pass the thresholds of p.adjusted < 0.1 and log2FC >1.

### Supplementary Figure S7

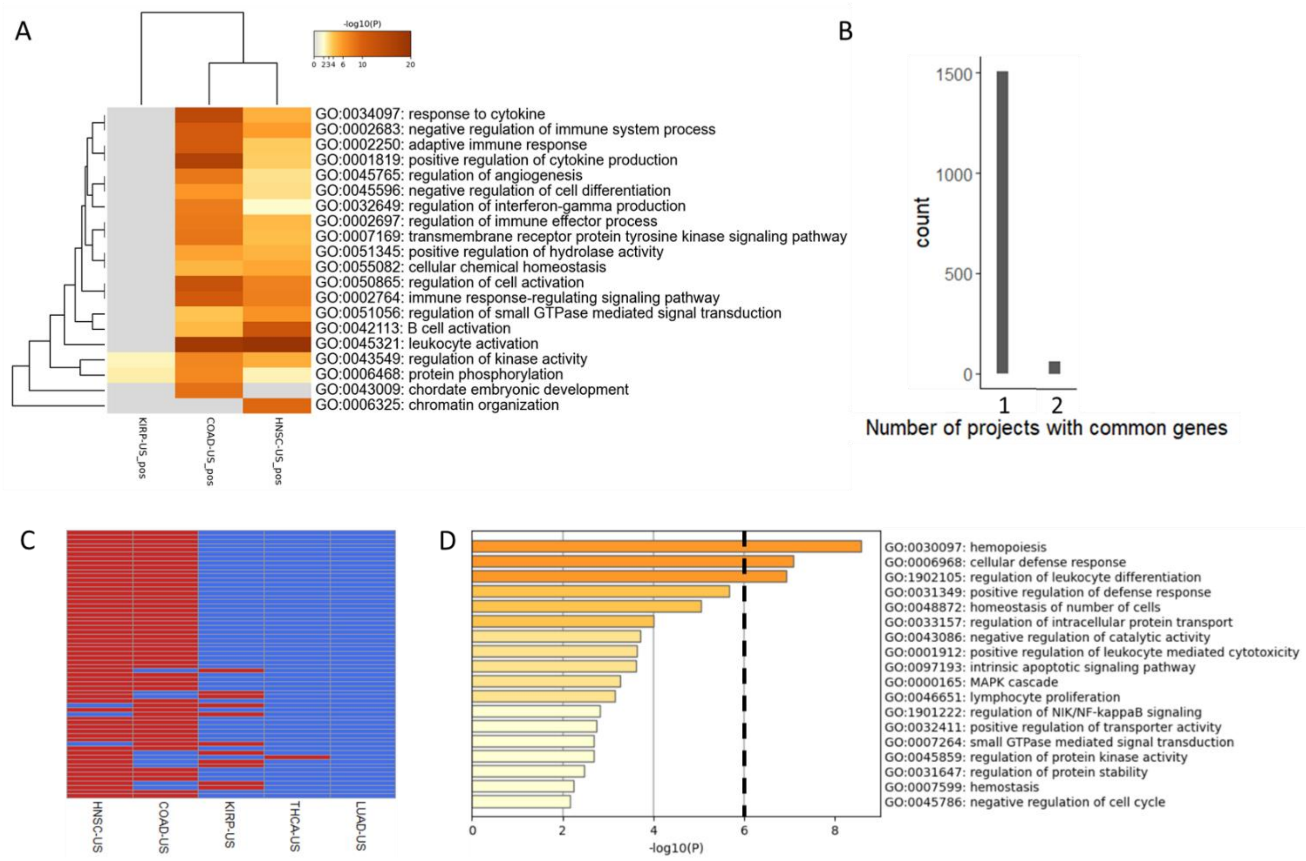


**Supplementary Figure S7. Expression profile analysis by Wilcoxon test of genes that expressed higher in the low RT-MR group**

Same as figure 4 B-E using ony genes differentially expressed by a Wicoxon test rather by Deseq2 statistics.

### Supplementary Figure S8

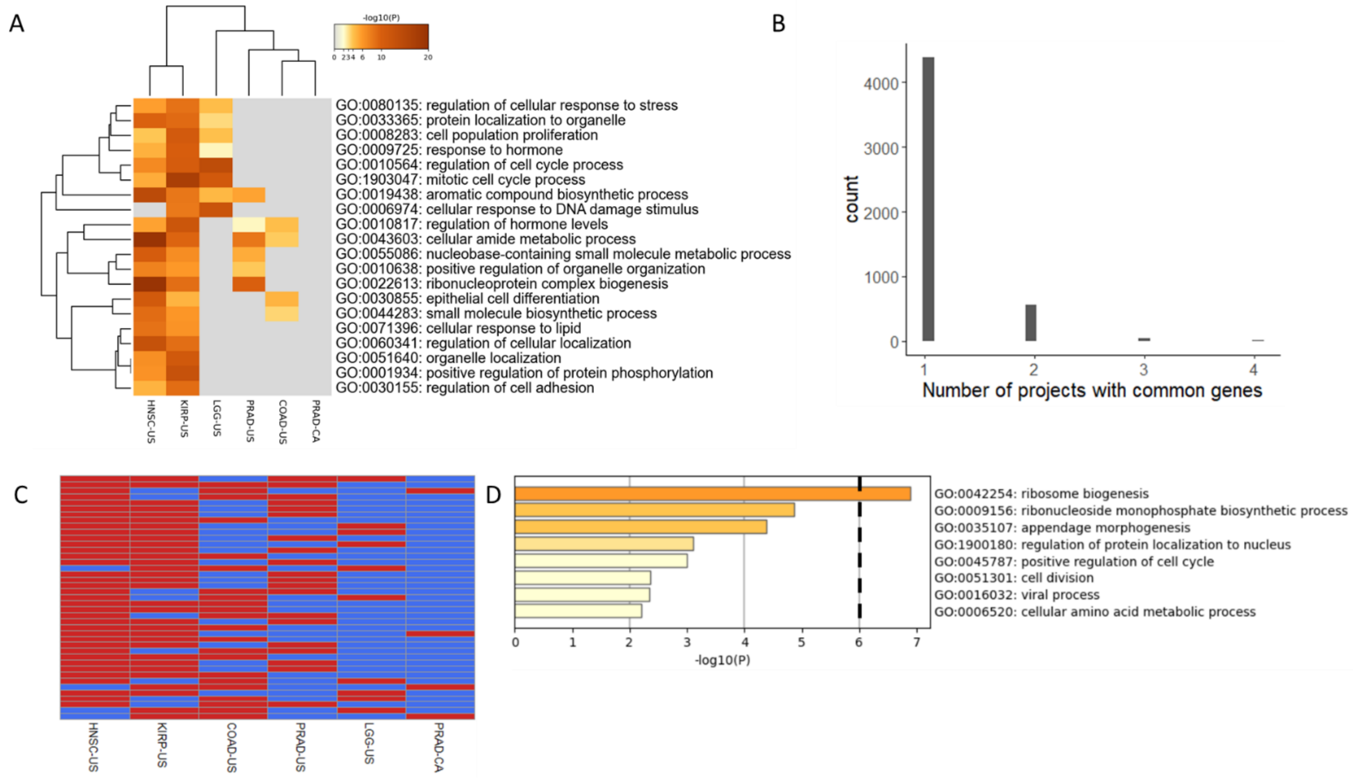


**Supplementary Figure S8. Expression profile analysis of genes that expressed higher in the high RT-MR group**

Same as figure 4 B-E for the complementary group of genes that are expressed higher in tumors with high RT-MR association.

### Supplementary Figure S9

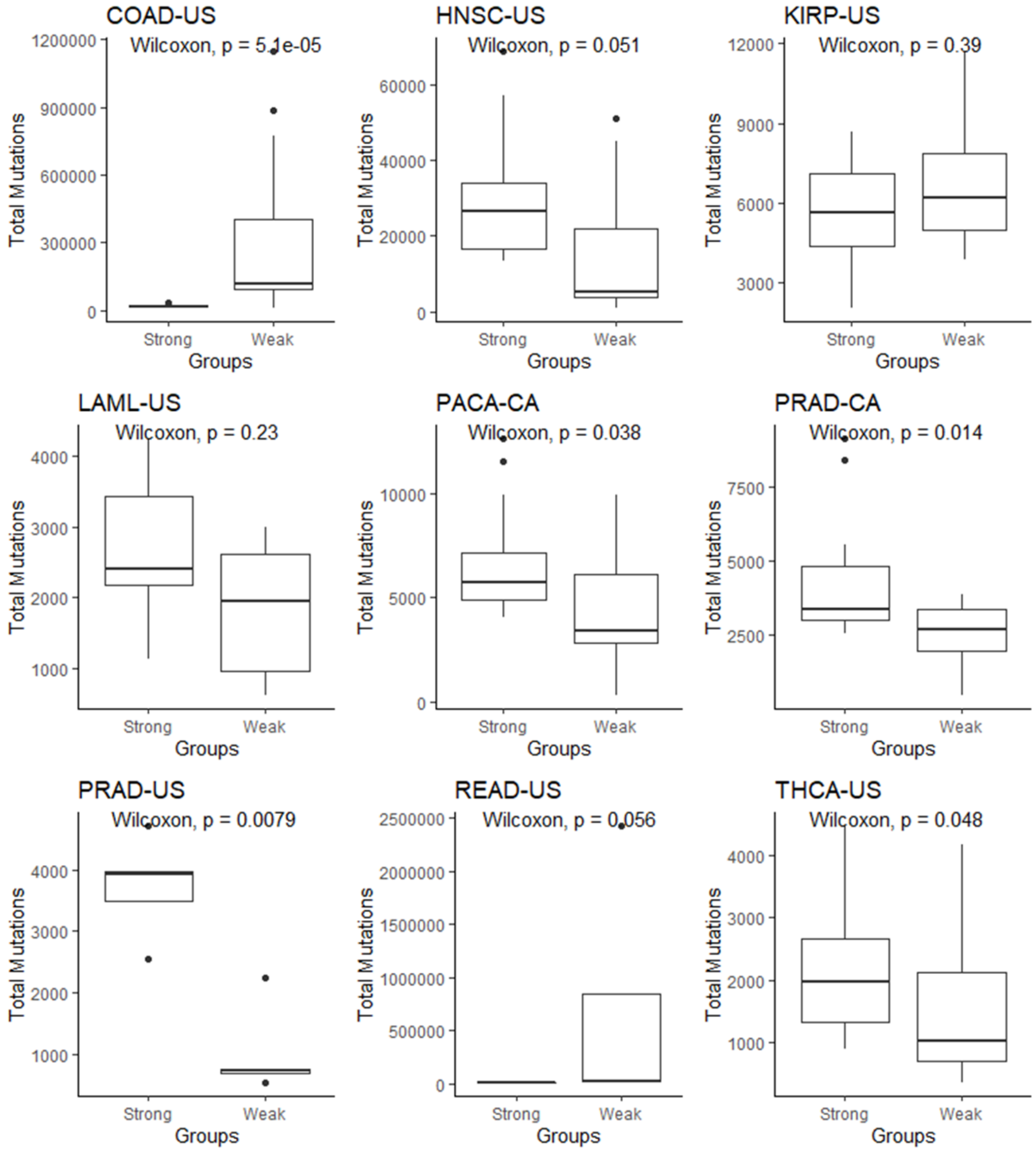


**Supplementary Figure S9. Total mutations of the strong and weak groups**

Boxplot of the total mutation's distribution of the strong and weak groups in the different projects. All P values derived from Wilcoxon rank–sum test.

### Supplementary Figure S10

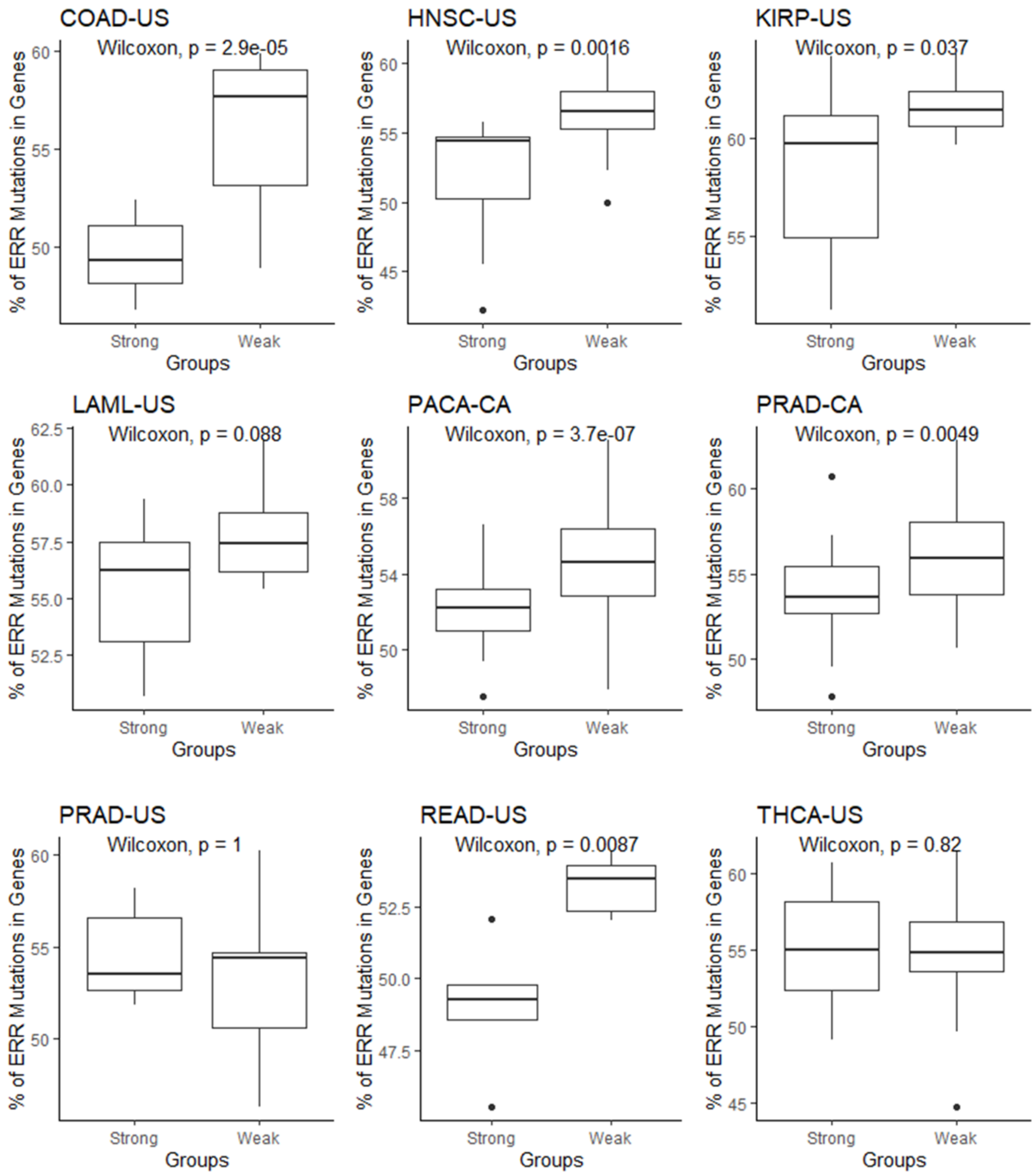


**Supplementary Figure S10. ERR mutations in the strong and weak groups**

Boxplot of the distribution of the percentage of ERR mutations in genes in the strong and weak groups in the different projects. The percentage was calculated by dividing the number of mutations that fell withing genes to the total number of mutations in early replicating regions in each sample. All P values derived from Wilcoxon rank–sum test.
